# Trans-Kingdom dsRNA Sensing: *Aspergillus fumigatus* Mycovirus Activates MDA5/MAVS Immunity and Limits Allergic Bronchopulmonary Aspergillosis

**DOI:** 10.64898/2026.05.18.725120

**Authors:** Alexander W. Rapp, Xi Wang, Brandon S. Ross, Alayna K. Caffrey-Carr, Sean M. Thomas, Agustin Resendiz-Sharpe, Andrew J. Olive, Katrien Lagrou, Robert A. Cramer, Joshua J. Obar

**Author notes:** Corresponding author: Joshua J. Obar, Geisel School of Medicine at Dartmouth, Department of Microbiology & Immunology, 1 Medical Center Drive, Lebanon, NH 03756. These authors contributed equally to this work (Alexander W. Rapp, Xi Wang).

## Abstract

MDA5 is a cytosolic pattern-recognition receptor (PRR) that binds to double-stranded RNA (dsRNA) and subsequently interacts with the signaling adaptor protein MAVS to initiate the antiviral interferon (IFN) response. Our group previously demonstrated that MDA5 is essential for host resistance against the fungal pathogen, *Aspergillus fumigatus*. Although fungal dsRNA was sufficient to activate MDA5 signaling, the precise source of *A. fumigatus* dsRNA responsible for this MDA5-stimulating function remains unknown. Here, we demonstrate that the magnitude of the IFN-dependent antifungal response is *A. fumigatus* strain dependent. Unexpectedly, we found that *A. fumigatus* isolates infected with dsRNA mycoviruses triggered a more robust MAVS-dependent inflammatory response within alveolar macrophages. Furthermore, dsRNA mycovirus infection increased fungal susceptibility to antifungal killing without altering other *A. fumigatus* growth characteristics. Although dsRNA mycovirus infection did not alter virulence in an acute bronchopneumonia model of *A. fumigatus* infection, it significantly impaired virulence and improved disease parameters in a chronic model of allergic bronchopulmonary aspergillosis (ABPA). Collectively, these findings reveal a novel role for trans-kingdom interactions in driving the host antifungal IFN response and modulating virulence in chronic aspergillosis models.

## Introduction

*Aspergillus fumigatus* is a ubiquitous environmental filamentous fungus capable of causing a spectrum of diseases in humans. The most severe form of disease is invasive aspergillosis, which predominantly affects neutropenic and highly immunosuppressed patients (1), but it is also emerging as a significant secondary infection following severe respiratory virus infections (2). In addition to these acute manifestations, *A. fumigatus* is also associated with several chronic disorders (3). Among these, allergic bronchopulmonary aspergillosis (ABPA) is a severe allergic disease driven by long-term *A. fumigatus* airway colonization, particularly in individuals with asthma and people with cystic fibrosis (pwCF). Although numerous murine models of *A. fumigatus-*driven allergic disease have been developed (4), until recently none adequately recapitulated the fungal persistence of ABPA (5). Consequently, the pathogenesis of ABPA and strategies to alleviate its symptoms remain incompletely understood (6).

Current first-line treatment for ABPA includes corticosteroids, such as prednisolone, and treatment with antifungal agents, such as the triazoles is sometimes utilized (7). In recent years, biologics like omalizumab (anti-IgE) (8), dupilumab (anti-IL4) (9), and anti-IL-5/IL-5R therapies (10) have also been used to manage uncontrolled ABPA, with variable success. To expand the therapeutic armamentarium for patients with ABPA, a deeper understanding of the host-fungal interactions driving this disease is needed. Of particular therapeutic interest is the potential use of mycoviruses, viruses that can induce hypovirulence in some pathogenic fungi (11). Additionally, understanding what fungal and host factors enable or limit *A. fumigatus* persistence to drive ABPA could aid in future treatment approaches.

Mycoviruses are viruses that exclusively infect fungi and are currently classified into 23 distinct families by the International Committee on the Taxonomy of Viruses (ICTV) (12). These viruses typically lack an extracellular phase of their life cycle and are transmitted vertically and/or horizontally (13). The majority of characterized mycoviruses, approximately 62%, possess a double-stranded RNA (dsRNA) genome and encode their own RNA-dependent RNA polymerase (RdRp) (14). Mycoviruses have been identified across all major fungal clades (15), and their impact on fungal virulence is complex and highly context-dependent. To date, most mycovirus research has focused on plant-associated fungal pathogens, in which hypovirulent, hypervirulent, and phenotypically neutral outcomes have all been reported with viral infected strains (16). Of relevance to human disease, mycoviruses have been identified in certain strains of *A. fumigatus* (17)*, Talaromyces marneffei* (18), and *Malassezia* spp.(19,20), where they can influence fungal virulence.

The impact of mycoviral infection on *A. fumigatus* virulence is complex and incompletely resolved (19); however, relatively little work has integrated these virulence studies with analyses of host immunity and chronic fungal disorders. We previously demonstrated that MDA5/MAVS signaling is essential for antifungal immunity against *A. fumigatus* in both mice and humans (20), (21). Additionally, type I and type III interferons (IFNs) have been demonstrated to be critical for antifungal immunity against *A. fumigatus* (48). Our previous work also established that dsRNA isolated from *A. fumigatus* was sufficient to trigger MDA5 activation and type I IFN expression (20), but the precise source of immunostimulatory dsRNA within the fungal RNA pool was not identified. Given that MDA5 sensing of dsRNA has been shown to promote fungal clearance (22) and the majority of characterized mycoviruses possess dsRNA genomes (15), we sought to determine whether mycoviral infection of *A. fumigatus* contributes to the antifungal type I interferon response observed in our prior studies. Here, we show that mycovirus infection of *A. fumigatus* is associated with enhanced type I IFN responses both *in vivo* and *in vitro,* and attributable to the presence of mycoviral dsRNA genomic segments. Mycoviral infection also increased *A. fumigatus* susceptibility to hydrogen peroxide and macrophage-mediated killing. As reactive oxygen species (ROS)-mediated killing is a major host defense mechanism against *A. fumigatus* (22), we further evaluated the consequences of mycoviral infection on *A. fumigatus* virulence across multiple disease models. Mycoviral infection reduced fungal burdens across all aspergillosis models tested; however, significant effects on disease outcomes were observed only in a chronic ABPA-like model.

## Methods

### Mice

*Mavs^fl/fl^* and *Mavs^fl/fl^* × *Itgax-Cre* mice were originally obtained from Dr. Sonja Best (23). *Ifih1^(-/-)^* mice were originally obtained from The Jackson Laboratory (Stock #015812). Transgenic mice were bred in-house at the Geisel School of Medicine at Dartmouth. C57BL/6J mice were purchased from The Jackson Laboratory (Stock #000664) and CD1 mice were purchased from Charles River (Strain #022). All mice were 8–16 weeks of age at the time of challenge. Mice were housed in autoclaved cages at ≤4 mice per cage with HEPA-filtered air and water. Animals were monitored daily for disease symptoms in accordance with our protocol. All animal studies were carried out in strict accordance with the recommendations in the *Guide for the Care and Use of Laboratory Animals* (24). All animal experiments were approved by the Institutional Animal Care and Use Committee (IACUC) at Dartmouth College (Protocol #00002168) or Michigan State University IACUC (animal use form [AUF] no. PROTO202200127).

### Fungal challenge models

For acute bronchopneumonia experiments, 8- to 12-week-old female C57BL/6J or CD1 mice were anesthetized by isoflurane inhalation and oropharyngeal challenged with 4×10^7^ live conidia in 100 μl PBS. Mice were sacrificed 48 hours post-challenge, as previously described (25).

For allergic bronchopulmonary aspergillosis (ABPA) experiments, 8- to 12-week-old female C57BL/6J mice were sensitized intranasally with 1×10^7^ live conidia on day -14, -7, -6, -5, -4, and -3, as previously described (5). Mice received a final intranasal challenge dose of 1 × 10^7^ conidia on day 0 and were sacrificed on day 7 post-challenge. All intranasal inoculations were performed under isoflurane anesthesia.

### *A. fumigatus* growth and collection of conidia

*A. fumigatus* strains used in this study are listed in Supplemental Table 1. Strains were grown on 1% glucose minimal medium (GMM) agar plates for 3 days at 37°C under normoxic conditions (21% oxygen, 5% CO_2_). Conidia were harvested by resuspension in 0.01% (v/v) Tween-80 in phosphate-buffered saline (PBS), filtered through sterile Miracloth, washed three times in sterile PBS, and quantified using a hemocytometer.

### Isolation of fungal RNA and screening *A. fumigatus* strains for mycoviral dsRNA

Fungal biofilms were established by inoculating freshly harvested conidia from 3-day-old GMM plates into liquid GMM (L-GMM) and culturing overnight 37°C with shaking under normoxic conditions. The following day, biofilms were lysed via a bead-beating in TRIzol^®^ Reagent for four 30-second intervals. Total RNA was then purified using a Zymo Research *Quick-*RNA Microprep Kit (Cat. #R1050) according to the manufacturer’s instructions.

For enrichment of dsRNA from the total RNA pool, 50 μg of fungal RNA was digested with RNase S1 (ThermoFisher, Cat. #EN021) at room temperature according to the manufacturer’s instructions. Residual genomic DNA was subsequently removed by digestion with DNase I (ThermoFisher Cat. #EN0521) at 37°C for 20 min. Reactions were terminated by the addition of 2 μl of 0.5M EDTA and enzymes were heat-inactivated at 75°C for 15 mins. Samples were immediately returned to ice to promote reanneal of dsRNA strands. Samples were then mixed with 3× formaldehyde loading dye (ThermoFisher Cat. #AM8552) and denatured at 65°C for 15 min on a heat block. dsRNA was resolved on a 2% agarose gel containing 1× NorthernMax^TM^ Denaturing Gel Buffer (Cat. #AM8676) and stained with SYBR^TM^ Green II RNA Stain (Cat. #S7564). Millenium^TM^ RNA markers (Cat. #AM7150) were used as a size reference.

### Screening *A. fumigatus* strains via qRT-PCR

RNA was isolated from fungal biofilms grown in L-GMM at 37°C with shaking, as described above, using a Zymo Research *Quick-*RNA Microprep Kit (Cat. #R1050). Complementary DNA (cDNA) was synthesized using a Qiagen QuantiTect Reverse Transcription Kit (Cat. #205311) according to the manufacturer’s instructions. Quantitative reverse-transcription PCR (RT-PCR) was performed using Af-PmV1-specific primers (Supplemental Table 2) and Bio-Rad SsoAdvanced Universal SYBR^®^ Green Supermix (Cat. #1725271) on a Bio-Rad CFX Opus 96 Real-Time PCR System, following the manufacturer’s recommended cycling conditions. The *A. fumigatus* translation elongation factor gene *tef1* (*Afu1g06390*) served as the endogenous reference gene, and relative viral gene expression was calculated using the ΔΔCt method.

### Generation of mycovirus-cured *A. fumigatus* isolates

Af-PmV1-infected strains were cured using previously published methods (26, 27).

For the cycloheximide curing approach, cycloheximide (MilliporeSigma, Cat. #C7698) was added to GMM agar plates at a final concentration of 150 μg/ml. One hundred conidia were spotted onto cycloheximide-containing plates, grown for 5 days at 37°C, and harvested by resuspension in PBS containing 0.01% (v/v) Tween-80 followed by filtration through Miracloth. This serial passaging process was repeated for a total of ten passages. Conidia were then diluted in PBS, spread onto GMM plates to obtain single colonies, and individual colonies were cultured overnight in L-GMM and screened for Af-PmV1 mycoviral genes using virus-specific primer sets (Supplemental Table 2).

For the ribavirin curing approach, one hundred spores were spotted onto GMM plates containing 0.1 mg/ml ribavirin (Cayman Chemical, Cat. #16757) and grown for 5 days at 37°C. of growth. Agar plugs were harvested, diluted, and plated onto GMM plates to obtain single colonies. Individual colonies were then screened by qRT-PCR for the presence of Af-PmV1 genes using mycovirus-specific primers (Supplemental Table 2).

For both approaches, all resulting isolates were additionally screened for residual dsRNA by treating total fungal RNA with RNase S1 and DNase and resolving remaining dsRNA by denaturing agarose gel electrophoresis, as described above.

### Re-infection of mycovirus-cured *Asfu1608r* isolate

Asfu1608r spores previously confirmed to be mycovirus-negative by qRT-PCR were re-infected via the transfection of Af-PmV1 dsRNA isolated from Asfu1608 back into Asfu1608r fungal protoplasts, as previously described (28). Briefly, spores were isolated from GMM agar plates and incubated at 28°C for 10-12 hours shaking in L-GMM to swell conidia to the germling stage. 50mg of lysing enzymes from *Trichoderma harzianum* was used to digest away the fungal cell wall by incubating at 28°C for roughly 5 hours. 10 μg of Asfu1608 dsRNA was transfected into Asfu1608r protoplasts following the removal of ssRNA from the total RNA pool as described above. Viable transformants were screened for the presence of Af-PmV1 dsRNA by resolving the dsRNA segments on a 2% agarose gel containing 1× NorthernMax^TM^ Denaturing Gel Buffer (Cat. #AM8676) and stained with SYBR^TM^ Green II RNA Stain (Cat. #S7564). Millenium^TM^ RNA markers (Cat. #AM7150) were used as a size reference. Additionally, qRT-PCR was used to screen for the presence of Af-PmV1 dsRNA segments.

### Genome sequencing and variant analyses

Mycelial cultures of *A. fumigatus* using L-GMM supplement with yeast extract were inoculated in small petri dishes grown overnight (18 to 24 h) at 37°C. Mycelia were collected, lyophilized, and bead beaten to powder, and DNA was extracted as previously described (29). Seacoast Genomics Inc. did genomic sequencing libraries and DNA sequencing. Analysis of impactful single nucleotide polymorphisms within the *A. fumigatus* genome utilized the Galaxy web platform (30) and SnpEff tools (31) as we have previously done (32).

### Stimulation of Primary Murine Fibroblast Cultures

Primary murine fibroblasts were isolated from ears of 6- to 12-week-old C57BL/6J mice. Ears were collected, washed in 70% ethanol, rinsed in PBS, and finely minced with scissors. Ear tissue from each mouse was then digested in 500 μl of 1,000 U/ml collagenase (Life Technologies, Cat. #17101-05) in Hank’s Balanced Salt Solution (HBSS) at 37°C for 25 min gently agitation. Digested tissue was centrifuged at 500 × *g* for 3 min, washed once with HBSS, and subsequently digested in 500 μl of 0.05% trypsin in HBSS for 20 mins at 37°C. Samples were then centrifuged again at 500 × *g* for 3 mins, the trypsin-containing supernatant was aspirated, and the tissue pellet was resuspended in 0.5 ml of fibroblast culture medium [DMEM supplemented with 10% fetal bovine serum (FBS), 1% MEM nonessential amino acids, and 1% penicillin-streptomycin]. Tissue was passed through a sterile 40-μm cell strainer, and the resulting cell suspension was used to initiate fibroblast cultures in tissue culture flasks with 25 ml of fibroblast culture medium. Medium was changed every 2 days, and fibroblasts were harvested at confluency (approximately day 6) using 0.25% trypsin.

For stimulation experiments, 50 ng of fungal dsRNA was packaged into LyoVec^TM^ liposomes per the manufacturer’s instructions and transfected into 5 × 10^4^ fibroblasts plated in 24-well plates. Following overnight incubation, cell supernatants were collected, and cytokine concentrations were measured by ELISA.

### Alveolar Macrophage (AlvMϕ)-like cell isolation, culture, and *in vitro* infection

Fetal liver–derived alveolar macrophage-like cells (FLAMs) were generated as previously described (33). Briefly, pregnant dams were euthanized by CO_2_ inhalation for 10 min, followed by cervical dislocation as a secondary method of euthanasia. Fetuses were immediately removed; loss of the maternal blood supply served as a method of fetal euthanasia. Cells were cultured in complete RPMI 1640 supplemented with 10% FBS and 1% penicillin-streptomycin. Where indicated, medium was further supplemented with 30 ng/ml recombinant mouse GM-CSF (PeproTech), and 15 ng/ml recombinant human TGF-β1 (PeproTech). Medium was refreshed every 2–3 days. When cells reached 70–90% confluency, they were detached by incubation for 10 min in cold PBS containing 10 mM EDTA, followed by gentle scraping.

To generate *Mavs*^(-/-)^ FLAMs, CRISPR/Cas9-mediated gene editing was employed. A *Mavs-*targeting single-guide RNA (sgRNA) (34) was cloned into the sgOpt lentiviral vector (Addgene plasmid no. 85681) (35). The resulting sgRNA construct was packaged into lentiviral particles as previously described and used to transduce early-passage Cas9-expressing FLAMs. Transduced cells were selected with puromycin beginning 2 days post-transduction.

For *in vitro* stimulation experiments, 5×10^5^ FLAMs were seeded into 24-well plates and challenged with live conidia at a multiplicity of infection (MOI) of 10 conidia per cell.

### Flow cytometry and Fluorescent Aspergillus Reporter (FLARE) analysis

Whole-lung single-cell suspensions were prepared and the fluorescent *Aspergillus* reporter strain AF293-FLARE was used as described previously (36). Briefly, lungs were minced and enzymatically digested in buffer containing 2.2 mg/ml collagenase type IV (Worthington), 100 μg/ml DNase I (Zymo Research), and 5% FBS at 37°C with rotation for 45 min. Digested tissue was passed through an 18-gauge needle, resuspended in red blood cell lysis buffer, diluted with PBS, filtered through a 100-μm mesh filter, and counted. For flow cytometric analysis, cells were stained with the antibody panel listed in Supplemental Table 3 and acquired on a Cytek^®^ Aurora Spectral Flow Cytometer in the DartLab Core Facility. Data were analyzed with FlowJo v10.7.1.

Cell populations were identified using the following gating strategy: alveolar macrophages (CD45^+^ MerTK^+^ CD64^+^ CD11b^(-)^ CD11c^+^ MHCII^+^ SiglecF^+^), interstitial macrophages (CD45^+^ MerTK^+^ CD64^+^ CD11b^+^ MHCII^+^), dendritic cells (CD45^+^ MerTK^(-)^ CD64^(-)^ CD11c^+^ MHCII^+^), monocytes (CD45^+^ MerTK^(-)^ CD64^(-)^ Ly6C^+^ CD11b^+^), neutrophils (CD45^+^ MerTK^(-)^ CD64^(-)^ SiglecF^(-)^ MHCII^+^ Ly6G^+^ CD11b^+^), and eosinophils (CD45^+^ MerTK^(-)^ CD11b^+^ CD64^(-)^SiglecF^+^ MHCII^(-)^).

### Cytokine analysis of BALF and blood serum from mice

At the indicated time-points after *A. fumigatus* challenge, mice were euthanized by CO_2_ inhalation. Bronchoalveolar lavage fluid (BALF) was collected by instilling 2 ml of PBS containing 0.05M EDTA through a cannula inserted into a tracheal incision. BALF was clarified by centrifugation at 300 × *g* for 5 min and stored at -20°C until analysis. BALF cytokine concentrations were measured by ELISA for CXCL10 (R&D Systems, Cat. #DY466), IFN-β (R&D Systems, Cat. #MIFNB0), IFN-α (Thermo Fisher Scientific, Cat. #BMS6027), IL-1α (BioLegend, Cat. #433401), IL-6 (BioLegend, Cat. #431316), TNF-α (BioLegend, Cat. #430916), IL-1β (BioLegend, Cat. #432601).

For serum collection, cardiac punctures were performed immediately after euthanasia, and blood was collected into serum separator tubes. Samples were centrifuged at 2,000 × *g* for 30 min, and serum was isolated from the supernatant layer. Total serum IgE was quantified by ELISA (Thermo Fisher Scientific, Cat. #EMIGHE) per the manufacturer’s instructions.

### Fungal burden determination

Culturable *A. fumigatus* was quantified from whole-lung homogenates. Lungs were homogenized in 1 ml sterile PBS using a Mini-BeadBeater (BioSpec Products) with glass beads. Serial dilutions (1:10 to 1:1,000) were plated onto Sabouraud dextrose agar and incubated overnight at 37°C. Colony-forming units (CFUs) were enumerated from plates containing 50-100 visible colonies and expressed as CFU per ml of lung homogenate.

For quantitative PCR (qPCR)-based fungal burden assessment, whole lungs were minced, freeze-dried overnight, and total DNA was extracted using an E.Z.N.A.^®^ Plant & Fungal DNA Kit (Omega Bio-Tek, Cat. #D3485-01). *Aspergillus*-specific primers targeting the 18S ribosomal DNA (rDNA) locus (Supplemental Table 2) were used to quantify total fungal burden by TaqMan^®^ Real-Time PCR analysis, as previously described (37,38).

### Histology: GMS and H&E analysis

For histological analysis, lungs were inflated with and fixed in 10% neutral buffered formalin for a minimum of 24 h. Fixed lungs were embedded in paraffin and sectioned at 5-μm thickness by the Dartmouth Health Pathology Shared Resource Laboratory (RRID: SCR_023479). Sections were stained with Grocott–Gömöri’s methenamine silver (GMS) to assess fungal germination and with hematoxylin and eosin (H&E) to evaluate lung inflammatory infiltrates, using standard histological techniques. Representative photomicrographs were acquired using an Olympus BX50WI microscope equipped with a QImaging Retiga 2000R digital camera.

### *In vitro* fungal peroxide killing assay

L-GMM was inoculated with either 2 × 10^6^ resting conidia or conidia pre-swollen in L-GMM at 37°C for 2 h. Both resting and pre-swollen conidia were then incubated at 37°C in L-GMM containing varying concentrations of hydrogen peroxide for 6 hours [AF293: 2mM H_2_O_2_; Asfu1608: 10mM H_2_O_2_]. Following incubation, serial dilutions were plated on GMM agar plates, and CFUs were enumerated and compared against PBS-only controls.

For experiments using the AF293-FLARE fluorescent reporter stain (36), both resting and preswollen conidia were exposed to a range of H_2_O_2_ concentrations (0-4 mM) prepared fresh in L-GMM. Loss of the RFP viability reporter signal was quantified by flow cytometry, and the percentage of dead conidia was calculated for both the Af-PmV1-infected and Af-PmV1-cured strains.

### Statistical Analysis

All statistical analyses were performed using GraphPad Prism software (v10.2.3). Unless otherwise indicated, data are presented as the mean ± standard error of the mean (SEM). For *in vitro* comparisons between two groups, two-tailed Student’s *t*-test was used. For animal experiments, parametric data from multiple groups were analyzed by one-way ANOVA followed by Tukey’s post hoc test; pairwise comparisons were made using Student’s *t*-test. For nonparametric data, the Kruskal-Wallis test with Dunn’s multiple comparisons correction was used for multiple-group comparisons, and the Mann–Whitney *U*-test was used for pairwise comparisons. Statistical significance thresholds are indicated as follows: NS, not significant; *, *P* ≤ 0.05; **, *P* ≤ 0.01; ***, *P* ≤ 0.001; ****, *P* ≤ 0.0001.

### Data availability

The genomic DNA sequences for the parental and mycovirus-cured strains are currently being deposited in NCBI GEO database.

## Results

### AF293 Induces an Enhanced IFN Response Compared to CEA10

Significant differences in the pro-inflammatory response and virulence across *A. fumigatus* strains has been noted (25,39), but how this fungal strain heterogeneity affects the fungal IFN response is ill-defined. Thus, we chose to directly compare the IFN-dependent inflammatory response of the AF293 and CEA10/A1163 reference strains of *A. fumigatus*. To assess whether the AF293 and CEA10 strains of *A. fumigatus* induce the interferon-dependent inflammatory response to different magnitudes, we challenged C57BL/6J mice with 4×10^7^ conidia oropharyngeally with either AF293 or CEA10. Forty-eight hours post-challenge, we examined cytokine levels in the bronchoalveolar lavage fluid (BALF) using a Luminex assay. When compared to AF293, we found that CEA10 induces significantly less CXCL9 and CXCL10 (Figure 1A-B). Importantly, this is not due to a global reduction in inflammation since CEA10 induced more IL-6 and IL-1α (Figure 1C-D) secretion into the BALF than AF293, which we have previously reported (25).

**Figure 1.**
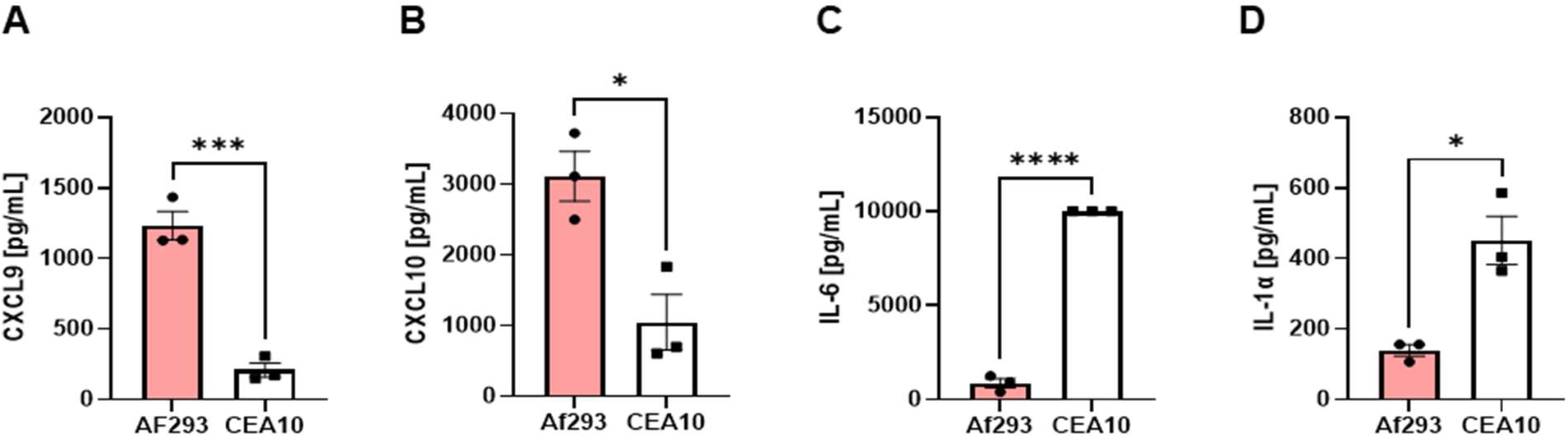
*Aspergillus fumigatus* strain AF293 elicits an enhanced type I IFN-mediated inflammatory response in the murine lung environment. **(A-D)** C57BL/6J mice (n = 3 per group) were intratracheally challenged with 4 × 10^7^ resting conidia of *A. fumigatus* strain AF293 or CEA10 in PBS. Bronchoalveolar lavage fluid (BALF) was collected 2 days post-challenge and analyzed for cytokine concentrations by ELISA. Each symbol represents an individual mouse; bars represent the mean ± one standard deviation (SD). Statistical significance was determined from one independent experiment using a two-way ANOVA (\**P* < 0.05;\*\*\**P* < 0.001;**** *P* < 0.0001).

### Alveolar Macrophages Induce More Potent Type I IFN Responses to AF293 than CEA10

Our previous results demonstrated that MAVS signaling in alveolar macrophages (AlvMϕ) was essential for host resistance and the induction of the type I and type III IFN-dependent inflammatory response (21). Moreover, we have demonstrated that fetal liver-derived AlvMϕ (FLAM), rather than bone marrow-derived macrophage are capable of inducing a robust type I IFN response to TLR2 agonists (40). Thus, we choose to use the AlvMϕ-like FLAM system to better understand this induction of this host-fungal interaction. Olive and colleagues demonstrate that both TGFβ and GM-CSF were necessary for the AlvMϕ-like phenotype of the FLAM cells, whereas GM-CSF only cells were more similar to monocyte-derived macrophages (33). Similar to our *in vivo* findings (21), AlvMϕ-like FLAM cells stimulated with AF293 conidia produced elevated levels of IFNβ and CXCL10 compared with GM-CSF only generated macrophages, but each produced equivalent levels of TNFα (Supplemental Figure 1). Additionally, AlvMϕ-like FLAM cells produced elevated levels of both IL-1α and IL-1β compared with GM-CSF only generated macrophages (Supplemental Figure 1). Thus, the AlvMϕ-like FLAM system is a good model for AlvMϕ interaction with *A. fumigatus*.

We next wanted to determine if a similar strain-specific inflammation response could be observed with the AlvMϕ-like FLAM cells. While the type I IFN-dependent inflammatory response was elevated in response to strain AF293, the pro-inflammatory cytokine response varied between strains (Figure 2A-E). As our previous work demonstrated that MAVS signaling in AlvMϕ was essential for host resistance against *A. fumigatus* (21), we next used our previously described CRISPR-Cas9 editing approaches in FLAMs to target *Mavs* (33). Similar to our earlier experiments, AF293 conidia induced greater IFNβ and CXCL10 secretion than CEA10 conidia in wild-type FLAMs. Moreover, following stimulation with AF293 conidia we observed decreased secretion of CXCL10 and IFNβ from *Mavs-*deficient FLAMs (Figure 2F-G). Thus, AF293 intrinsically leads to greater MAVS-dependent IFN response than CEA10.

**Figure 2.**
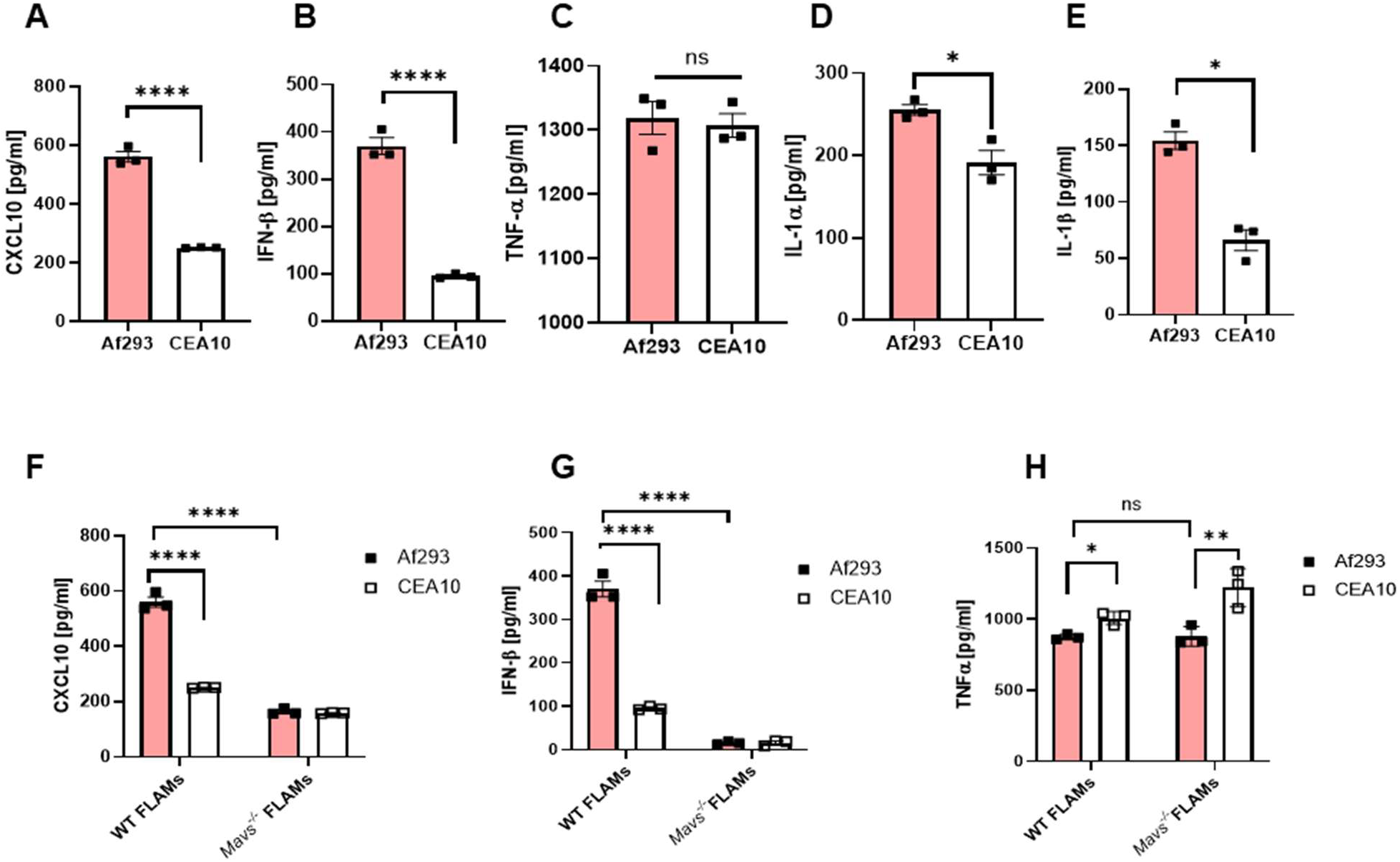
dsRNA mycoviral infection of *A. fumigatus* correlates with the strain-specific induction of a MAVS-dependent type I IFN response in alveolar macrophages. **(A-E)** FLAMs cultured with TGFβ and GM-CSF were challenged overnight with resting conidia of AF293 or CEA10 at an MOI of 10:1. CXCL10 and IFN-β concentrations were quantified by ELISA. Each symbol represents a single well; bars represent the mean ± SD. **(F-H)** Wild-type (WT) and *Mavs*^-/-^ FLAMs cultured with TGFβ and GM-CSF were challenged overnight with resting conidia of AF293 or CEA10 at an MOI of 10:1. CXCL10 and IFN-β concentrations were quantified by ELISA. Each symbol represents a single well; bars represent the mean ± SD. Statistical significance was determined from three independent experiments using a two-way ANOVA (ns = not significant; \**P* < 0.05; \**P* < 0.01; **** *P* < 0.0001).

### Af-PmV1 Mycoviral Infection Correlates with an Enhanced IFN-dependent Inflammatory Response

MAVS is a critical scaffolding and signaling adaptor protein for the RIG-I-like receptors MDA5 and RIG-I, which function as cytosolic RNA sensors (41). Our previous work established that MDA5 is essential for host resistance and the induction of the type I and type III IFN-dependent inflammatory response following *A. fumigatus* challenge (20). We also demonstrated that dsRNA isolated from AF293 was sufficient to trigger an type I IFN-dependent inflammatory response in murine fibroblasts (20). To directly compare the immunostimulatory properties of fungal RNA we initially purified total RNA from the overnight biofilms of AF293 and CEA10 reference strains that was then packaged into LyoVec™ liposomes and transfected into primary murine fibroblasts. Only RNA isolated from the AF293 strain stimulated IFNα, IFNβ and CXCL10 secretion (Figure 3A). We next extended this analysis to additional clinical isolates, identifying Asfu1608 as a potent inducer of the type I IFN response (Figure 3B). Our prior demonstration that the treatment of AF293 RNA with RNase III (specific for dsRNA) abrogates IFN-dependent inflammatory responses (20) suggests that strain-specific differences in IFN induction may reflect differential abundance of dsRNA within the total fungal RNA pool capable of driving robust MDA5 activation remains unresolved.

**Figure 3.**
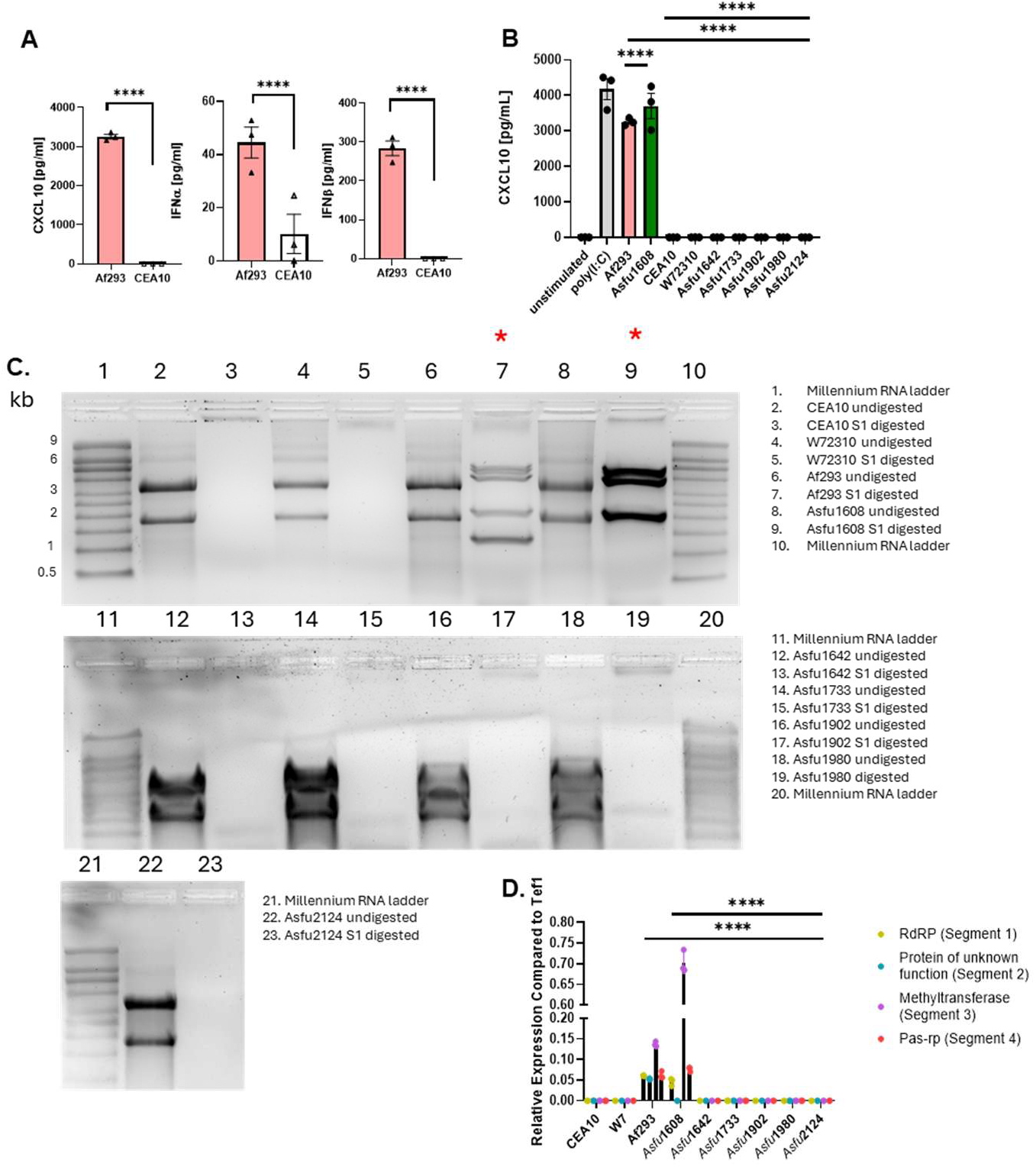
Af-PmV1 dsRNA is present exclusively in *A. fumigatus* strains that potently induce type I IFN responses. **(A-B)** A total of 0.05 μg of fungal RNA from overnight biofilms was encapsulated in LyoVec^TM^ liposomes and transfected into 5 × 10^4^ primary C57BL/6J murine fibroblasts. RNA from laboratory strains and clinical isolates was tested. Culture supernatants from overnight incubations were analyzed by ELISA for IFN-α and IFN-β production. Data are from a single experiment. Each triangle represents a single well, and the bar is the mean ± SD. Statistical significance was determined using either a Student’s *t*-test (**A**) or two-way ANOVA (**B**) (\*\*\*\**P* < 0.0001). (**C**) Total RNA from overnight biofilms was enriched from the same strains used in (**B**). Total fungal RNA was treatment with RNase S1 and DNase I to get rid of ssRNA and DNA, respectively. The remaining dsRNA was run on a 2% denaturing Northernmax^TM^ agarose gel. Red asterisks indicate strains where dsRNA was visualized following ssRNA digestion. (**D**) qRT-PCR was performed on cDNA made from bulk fungal RNA preps, testing for the presence of PmV1 viral genes. Relative expression of the viral genes was compared relative to the fungal translation elongation factor gene *tef1* (*Afu1g06390).* Statistical significance was determined from three independent experiments using a two-way ANOVA (\*\*\*\**P* < 0.0001).

Given that MDA5 was originally identified as a cytosolic receptor for viral dsRNA (42), we investigated whether the observed strain-specific induction of the type I IFN-dependent inflammatory response was attributable to mycovirus infection. Importantly, AF293 has previously been reported to harbor *Aspergillus fumigatus* polymycovirus 1 (Af-PmV1) (27, 44). To determine whether mycoviral infection was present in our panel of *A. fumigatus* strains, total RNA was extracted from overnight biofilm cultures, treated with DNase and RNase S1 to enrich for dsRNA, and resolved on denaturing agarose gels. This analysis revealed that only AF293 and Asfu1608, that both induce a strong type I IFN response (Figure 3A-B), contained abundant dsRNA bands, consistent with mycoviral infection (Figure 3C). In our AF293 stock, five dsRNA segments were visualized, suggesting infection with the Af*-*PmV1M variant as first described by Takahashi-Nakaguchi et al. (43), whereas Asfu1608 exhibited four dsRNA segments, consistent with the original Af-PmV1 description (44). To confirm mycoviral identity, we screened for Af*-*PmV1 open reading frames (ORFs), across four gene segments, by qRT-PCR using mycovirus-specific primers. This analysis confirmed that both AF293 and Asfu1608 were infected with Af*-*PmV1 or Af-PmV1M (Figure 3D).

### Af-PmV1 dsRNA Drives a type I IFN Response *In Vitro* and *In Vivo*

The data presented thus far are consistent with the hypothesis that Af-PmV1 infection contributes to the enhanced type I IFN-dependent inflammatory response observed in AF293 and Asfu1608. To directly test this hypothesis, we cured the AF293 and Asfu1608 strains of their mycoviral infections by one of two approaches: serial passaging on GMM plates containing cycloheximide (45), or culture on GMM plates containing ribavirin (26) (Figure 4A). The ribavirin-based curing method successfully cured both AF293 and Asfu1608 of their Af-PmV1 infection, whereas cycloheximide treatment was effective only for the AF293 strain (Figure 4C-D, Supplemental Figure 2). To confirm that the antiviral curing procedures did not introduce confounding genomic alterations, we utilized whole-genome sequencing on all parental and mycovirus-cured strains. This analysis identified 15 single-nucleotide polymorphisms (SNPs) in 15 distinct protein-coding sequences in AF293-dervived isolates following serial passaging on cycloheximide plates, four of which were shared between the cured (AF293c) and uncured (P10.2) isolates. For ribavirin-treated AF293 isolates, 20 SNPs were identified across both the cured (AF293r) and uncured (AF293rib-uncured) strains, with two SNPs shared between the cured and uncured isolates (Supplemental Table 4A). An equivalent analysis was performed for ribavirin-treated Asfu1608 cured and uncured isolates (Supplemental Table 4B). To rule out growth-related phenotypic differences as confounders of the differential IFN response, radial growth on GMM agar plates and conidial germination rates were compared between virus-infected and virus-cured isogenic strain pairs; no significant differences were observed (Supplemental Figure 3C-D).

**Figure 4.**
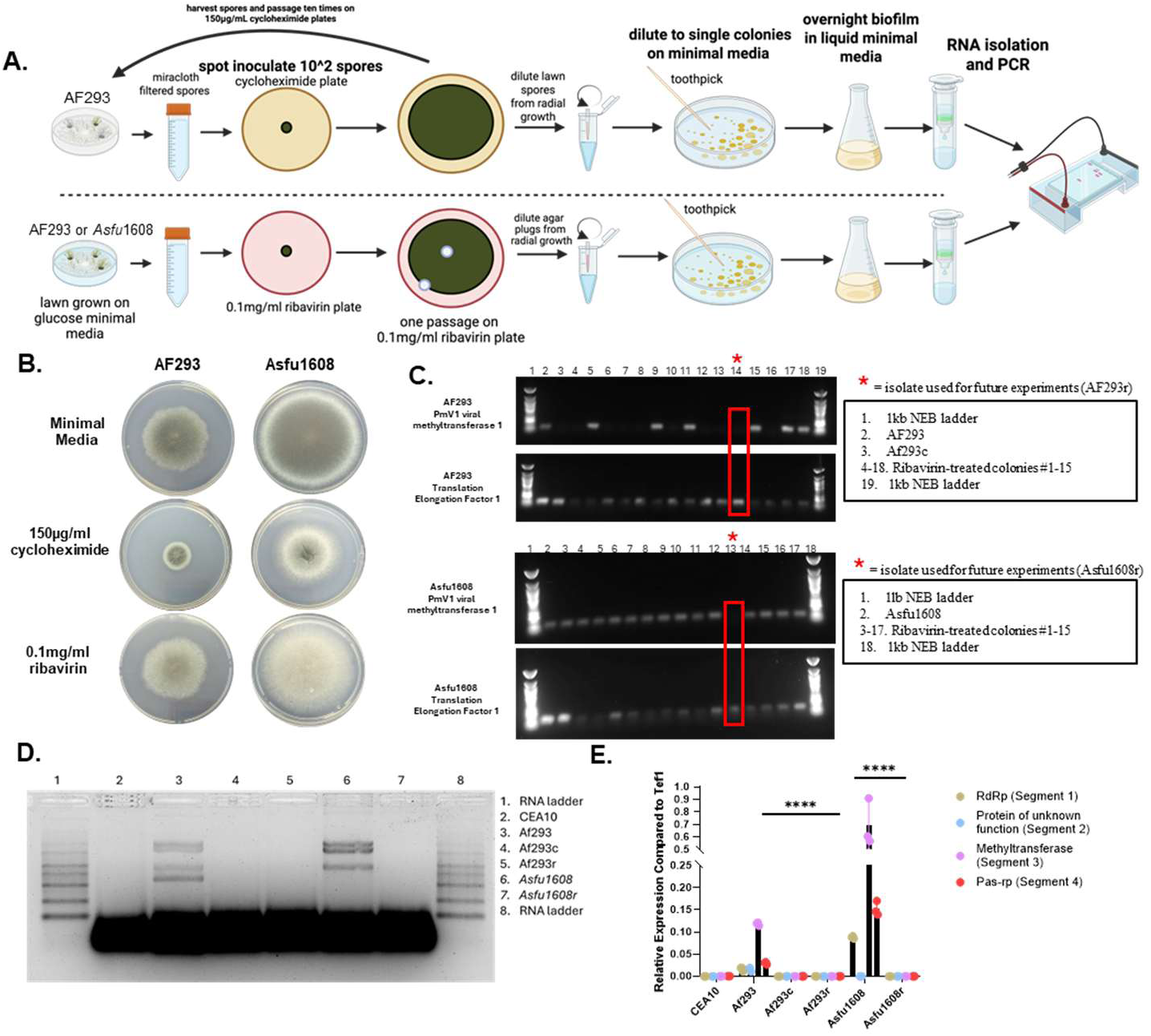
Ribavirin and cycloheximide treatment effectively cure *A. fumigatus* of Af-PmV1 infeciton without altering fungal growth characteristics, generating isogenic mycovirus-infection and mycovirus-free strain pairs. Strains AF293 and Asfu1608 were independently cured of their PmV1 viral infection via two independent methods. (**A**) Schematic diagram for mycovirus curing processes, in which 100 conidia of strain AF293 were spotted onto a GMM plate containing either 150 μg/ml cycloheximide or 0.1 mg/ml ribavirin. After 5 days of radial growth, resting spores were harvested and passaged on cycloheximide plates again for a total of ten passages. For AF293 and Afu1608 grown on ribavirin plates, agar plugs were taken from the pate and diluted to single colonies on a GMM plate. Single colonies were grown separately as a biofilm in L-GMM. Colonies were subsequently tested for the presence of Af-PmV1 genes via RT-PCR. Schematic was made at Biorender.com with appropriate licenses. (**B**) Growth phenotypes upon exposure to either ribavirin or cycloheximide after 5 days of growth. **(C)** PCR testing for the absence or presence of the Af-PmV1 methyltransferase gene following antiviral treatment. Fungal translation elongation factor gene *tef1* (*Afu1g06390)* used as a control. **(D)** dsRNA from a biofilm of each strain grown overnight was run on a 2% denaturing Northernmax^TM^ agarose gel to test for the complete absence of dsRNA in the cured strains. **(E)** qRT-PCR testing for the absence of viral genes following antiviral treatment of the PmV1+ strains. Relative expression of the viral genes was compared relative to the fungal translation elongation factor gene (*Afu1g06390) tef1.* Statistical significance was determined using a two-way ANOVA (\*\*\*\**P* < 0.0001).

With these isogenic mycovirus-infected and mycovirus-cured strains in hand, we could now directly evaluate the role of Af-PmV1 in the induction of the type I IFN-dependent inflammatory response. Initially, we purified total fungal RNA from the AF293, AF293c, and AF293r strains, packaged the fungal RNA into LyoVec™ liposomes, and then stimulated C57BL/6J fibroblasts. As we have previously observed, RNA isolated from AF293 was able to induce robust secretion of IFNβ and CXCL10, while RNA isolated from either of the mycovirus-cured strains (AF293c or AF293r) was severely diminished in its ability to stimulate IFNβ and CXCL10 production (Figure 5A). Similarly, RNA isolated from Asfu1608 was immunostimulatory, while RNA isolated from Asfu1608r was not (Figure 5A). Thus, the RNA genomic segments of Af-PmV1 are necessary for the immunostimulatory function of the fungal RNA pool isolated from AF293 and Asfu1608.

**Figure 5.**
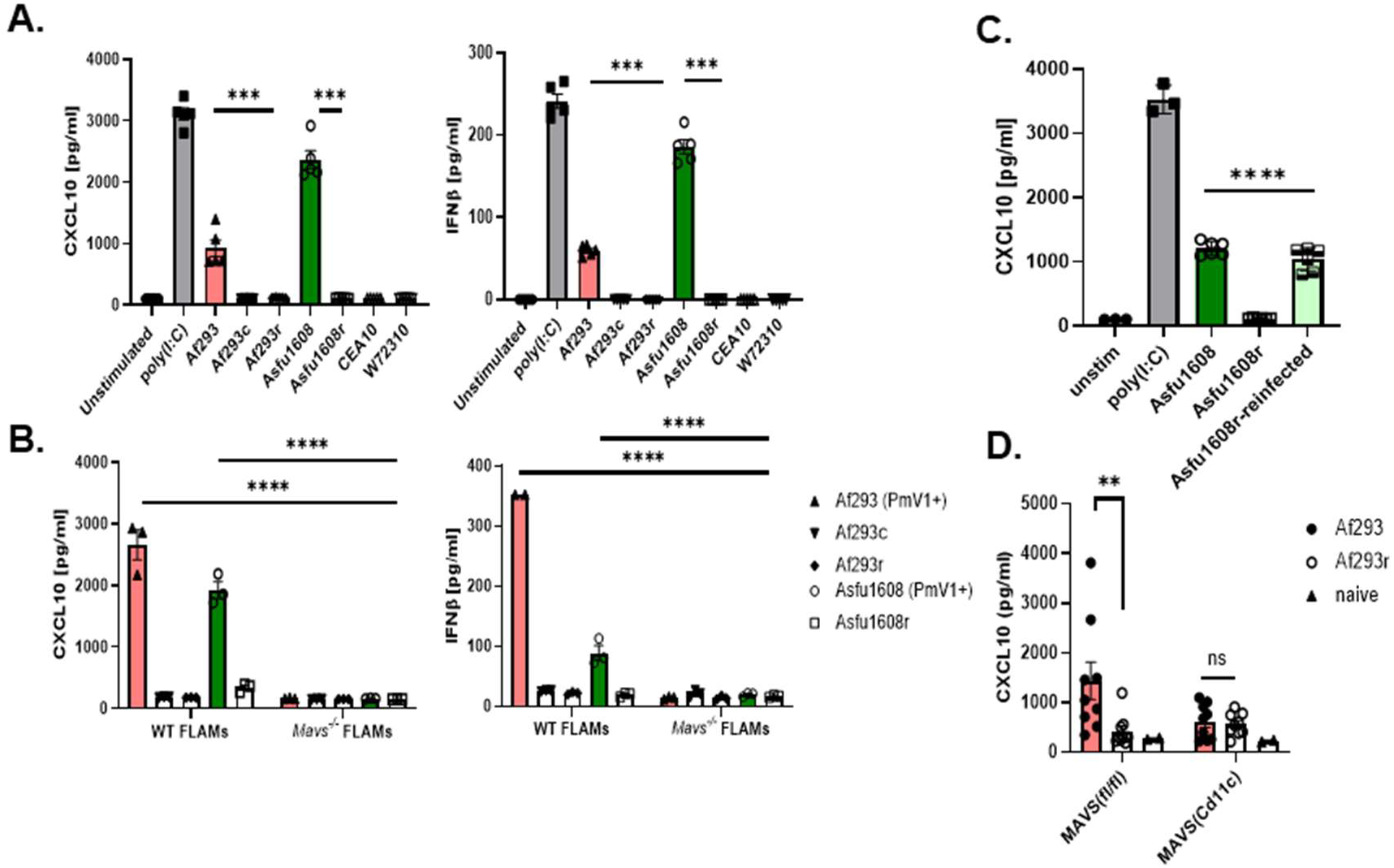
Mycovirus curing abrogates Af-PmV1-driven MAVS-dependent type I IFN responses in alveolar macrophages *in vitro* and *in vivo*. **(A)** 0.05μg of fungal RNA from overnight biofilms was packaged into LyoVec^TM^ liposomes and transfected into primary C57BL/6 primary mouse fibroblasts. Cell supernatant from overnight cell culture was analyzed via ELISA for production of interferons as in Figure 2. Each shape represents a single well, and the bar represents the mean ± one standard deviation. Statistical significance was determined using a two-way ANOVA (\*\*\**P* < 0.001; **** *P* < 0.0001). **(B)** WT and *Mavs^-/-^* FLAMs were cultured in the presence of 15ng/ml TGFβ were infected at an MOI of 10:1 with live spores of either PmV1 infected or cured strains of AF293 or Asfu1608. Following overnight infection, culture supernatants were analyzed for differences in CXCL10 and IFNβ production. Each shape represents a single well, and the bar represents the mean ± one standard deviation. Statistical significance was determined using a two-way ANOVA (\*\*\**P* < 0.001; **** *P* < 0.0001). **(C)** WT FLAMs were infected at an MOI of 10:1 with live spores of either PmV1 infected, cured, or re-infected strains of Asfu1608. Following overnight infection, culture supernatants were analyzed for differences in CXCL10 production. Each shape represents a single well, and the bar represents the mean ± one standard deviation. Statistical significance was determined using a two-way ANOVA (\*\*\*\**P* < 0.0001). **(D)** CD11c-cre x *Mavs^fl/fl^*mice and littermate controls were intratracheally infected with live spores of either PmV1 infected AF293 or AF293 cured with Ribavirin (AF293r). CXCL10 was quantified from the bronchoalveolar lavage fluid. Each shape represents a single mouse, and the bar represents the mean ± one standard deviation. Statistical significance was determined from a representative experiment using a two-way ANOVA (ns = non-significant; ** *P* < 0.001).

We next wanted to test whether the Af-PmV1 influences immunostimulatory properties of live *A. fumigatus* conidia. To assess the role of Af-PmV1 in triggering the type I IFN-dependent inflammatory response in AlvMϕ, we challenged wild-type and *Mavs-*deficient FLAMs with live resting conidia of the AF293/AF293c or Asfu1608/Asfu1608r isogeneic pairs of *A. fumigatus* strains. Our previous data demonstrated that AlvMϕ are the major hematopoietic cell type responsible for the MAVS-dependent sensing of *A. fumigatus* (21), and consistent with this, only of mycovirus-infected conidia (AF293 and Asfu1608), but not their mycovirus-cured derivatives (AF293c and Asfu1608r), were able to stimulate IFNβ and CXCL10 secretion from wild-type FLAMs; this response was abrogated in *Mavs-*deficient FLAMs (Figure 5B). Although the use of isogenic mycovirus-infected and mycovirus-free strain pairs strongly supports a causal role for mycoviral infection in contributing to the MAVS-dependent type I IFN response, antiviral curing procedures can introduce off-target genomic changes that could, in principle, confound this interpretation, a concern that our SNP analysis only partially addresses. To directly and definitively establish that Af-PmV1 infection is both necessary and sufficient for enhanced type I IFN response, we performed a viral reconstitution experiment in which Af-PmV1 infection was re-established in the mycovirus-cured Asfu1608r strain by transfecting purified Asfu1608 dsRNA into Asfu1608r protoplast (Supplemental Figure 2C-D). Strikingly, re-infection of Asfu1608r with Af-PmV1 fully restored the enhanced type I IFN response to levels comparable to the parental mycovirus-infected Asfu1608 strain, as measured by CXCL10 secretion from wild-type FLAMs (Figure 5C). Together, the loss-of-function (curing) and reconstitution (re-infection) data establish the Af-PmV1 infection is necessary and sufficient to contribute to the enhanced MAVS-dependent type I IFN response in AlvMϕ-like FLAM cells.

To extend these findings *in vivo,* we next assessed the role of Af-PmV1 in driving the IFN-dependent inflammatory response in the respiratory tract using mice with conditional deletion of *Mavs* in CD11c-expressing cells. *Mavs^(fl/fl)^* and *Mavs^(CD11c)^* mice were challenged i.t. with 4×10^7^ conidia of AF293 or AF293r, and CXCL10 levels in the BALF were quantified 48 hours post-challenge. In *Mavs^(fl/fl)^* mice, AF293 induced significantly greater CXCL10 than AF293r confirming that Af-PmV1 infection enhances the *in vivo* IFN response (Figure 5D). Importantly, in *Mavs^(CD11c)^* mice both AF293 and AF293r conidia induced comparable and similarly reduced levels of CXCL10, which was equivalent to the response AF293r conidia induced in *Mavs^(fl/fl)^* mice (Figure 5D). This demonstrates that the Af*-*PmV1 infection was responsible for the enhanced AlvMϕ- and MAVS-driven IFN-dependent inflammatory response following *A. fumigatus* challenge. Collectively, these data establish that AlvMϕ sense the Af-PmV1 dsRNA genome through a MAVS-dependent pathway to promote an enhanced type I IFN response during *A. fumigatus* infection.

### Af-PmV1 Infection Enhances Fungal Susceptibility to H_2_O_2_ and Leukocyte-mediated Killing

Mycovirus infection can produce hypovirulent, hypervirulent, or phenotypically neutral outcomes depending on the host-pathogen system under study (15). We therefore evaluated whether Af-PmV1 infection altered the ability of *A. fumigatus* to withstand host-relevant stresses. Because ROS-mediated killing represents the predominant leukocyte host defense mechanism against *A. fumigatus* (46), we first employed an *in vitro* hydrogen peroxide (H_2_O_2_) killing assay in which freshly harvested conidia were incubated with varying concentrations of H_2_O_2_ in liquid GMM at 37°C for 5 hours. Quantification of colony-forming units (CFUs) revealed that Af-PmV1 infected AF293 resting conidia were more susceptible to H_2_O_2_-mediated killing than the cured strain AF293c; however, pre-swollen Af-PmV1-infected AF293 conidia were conversely more resistant to H_2_O_2_-mediated killing (Figure 6A). This bidirectional, germination state-dependent effect was not observed for the Asfu1608 clinical isolate: Af-PmV1-infected Asfu1608 conidia were more susceptible to H_2_O_2_-mediated killing than the virus-cured strain Asfu1608r, but only in the pre-swollen state (Figure 6A). To establish that this enhanced H₂O₂ susceptibility is directly attributable to Af-PmV1 infection rather than off-target effects of antiviral curing, we again used the re-infected Asfu1608r strain. Strikingly, re-infection of Asfu1608r with Af-PmV1 restored H_2_O_2_ susceptibility to levels comparable to those of the parental Asfu1608 strain (Figure 6A), establishing that Af-PmV1 infection is both necessary and sufficient for the enhanced susceptibility to oxidative killing observed in this isolate.

**Figure 6.**
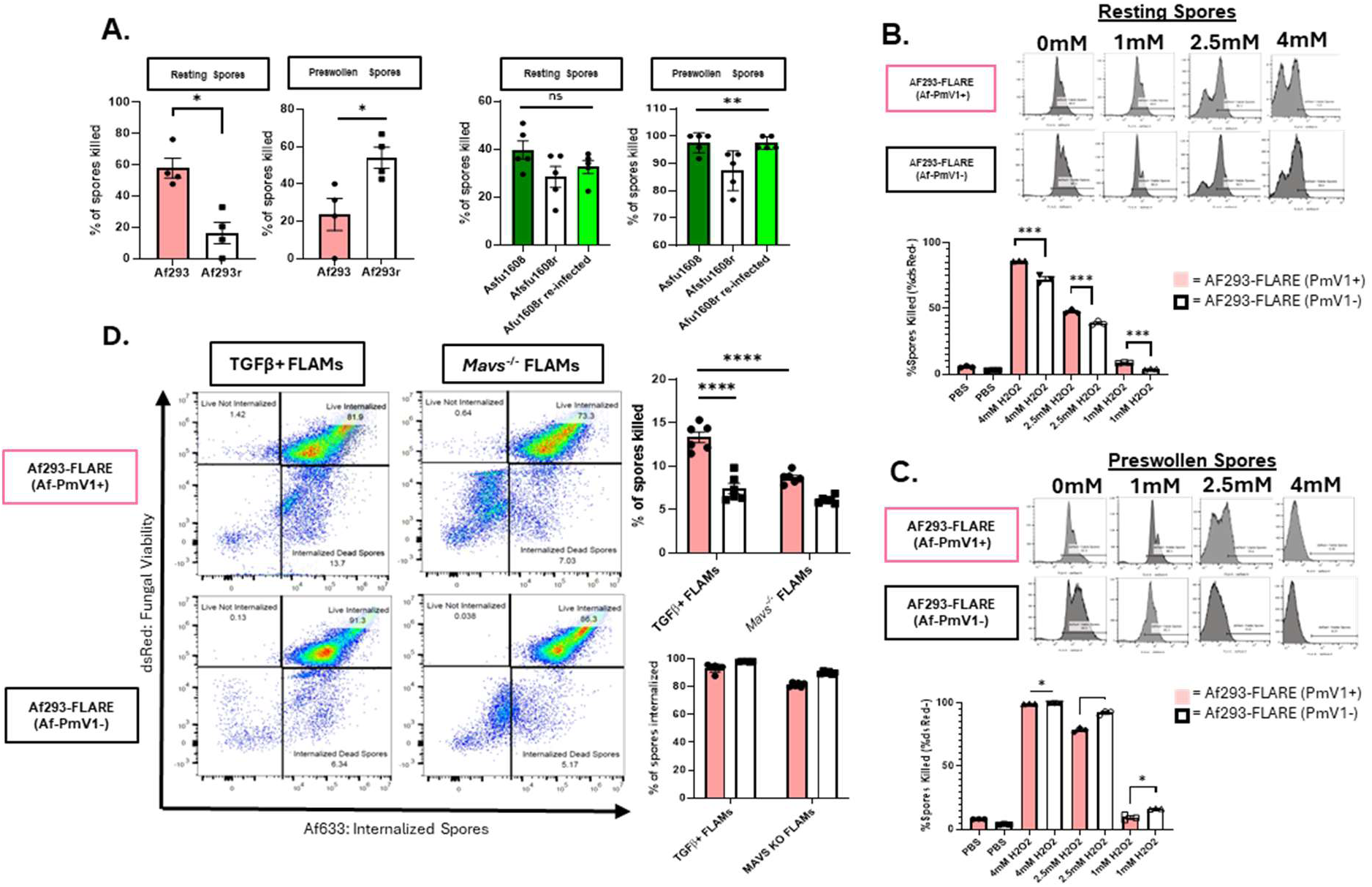
Af-PmV1 infection enhances *A. fumigatus* conidial susceptibility to H_2_O_2_- and alveolar macrophage-mediated killing in a germination state-dependent manner. **(A)**. Both resting and spores pre-swollen in L-GMM shaking at 37°C for 2 hours were incubated in varying concentrations of hydrogen peroxide for 6 hours at 37°C. AF293 was incubated with 2mM H_2_O_2_ while Asfu1608 was incubated with 10mM H_2_O_2_. CFUs were performed on the conidia following incubation and quantified in comparison to PBS only controls. (**B,C**) Resting and pre-swollen fluorescent viability reporter strains of AF293 (AF293-FLARE) were incubated in the presence of varying concentrations of hydrogen peroxide for 30 minutes at 37°C in L-GMM. Loss of the dsRed viability fluorescent protein was used as an indicator for spore death. Each dot represents a single well, and the bar represents the mean ± SD. Statistical significance was determined using a Student’s t-test from three independent experiments (ns = not significant, (* *P* < 0.05;** *P* < 0.01; *** *P* < 0.001; **** *P* < 0.0001). **(D)** 5 × 10^5^ wild-type and *Mavs^-/-^* FLAMs were infected with either the Af-PmV1 infected or ribavirin-cured fluorescent reporter strains of AF293 at an MOI 10 spores every 1 cell and incubated at 37°C for 10 hours.

To further validate that Af-PmV1 infection altered fungal susceptibility to antifungal immune cell-mediated killing and identify which host immune cell population is responsible for the enhanced clearance seen in the Af-PmV1-infected strain we utilized the *A. fumigatus* fluorescent reporter (FLARE) assay (36). To do this, we generated isogenic Af-PmV1-infected or Af-PmV1-free AF293 FLARE strains using the ribavirin curing method (Supplemental Figure 4) and validated in our *in vitro* peroxide exposure assay. Consistent with the CFU-based data (Figure 6A), resting Af-PmV1-infected AF293 conidia were more readily killed by H_2_O_2_ exposure, whereas pre-swollen conidia showed the reverse effect (Figure 6B-C). When we examined phagocytosis and killing of AF293 and AF293r conidia by AlvMϕ-like cells in an *in vitro* FLARE assay, both strains were phagocytosed at comparable levels; however, mycovirus-infected AF293 conidia were killed significantly more efficiently than their mycovirus-cured AF293r counterparts (Figure 6D). Furthermore, both AF293 and AF293r conidia were killed at comparable and reduced levels in *Mavs-*deficient AlvMϕ-like cells, which was comparable to the killing of AF293r conidia by wild-type AlvMϕ-like cells (Figure 6D). This demonstrates that Af-PmV1 infection contributes to enhanced AlvMϕ-mediated killing through a MAVS-dependent mechanism.

To extend these findings *in vivo*, we challenged C57BL/6J mice i.t. with 4×10^7^ resting conidia of either the Af-PmV1-infected and Af-PmV1-free AF293 FLARE strain and analyzed fungal uptake and killing by CD45^+^ immune cell populations within the lung parenchyma at 36 hours post-challenge, as has previously been done (36). Both alveolar macrophages and interstitial macrophages displayed significantly enhanced internalization and killing of Af-PmV1 infected conidia compared to mycovirus-free conidia (Figure 7B). In contrast, neutrophils exhibited a modestly decreased ability to kill Af-PmV1 infected conidia (Figure 7B). To quantify the net effect of these differential killing phenotypes on overall fungal clearance, immunocompetent C57BL/6J mice were challenged intranasally (1×10^6^ conidia) or oropharyngeally (4×10^7^ conidia) with either the pairs of AF293/AF293r or Asfu1608/Asfu1608r conidia. Lung fungal burdens were quantified by CFU assay at 48 hours post-challenge. Regardless of the route of administration, inoculum dose, or fungal strain background, Af-PmV1 infection was associated with significantly lower fungal burdens (Figure 7C-D). To establish that the reduced fungal burden associated with Af-PmV1 infection is directly caused by mycoviral infection rather than off-target curing effects, we again employed a viral reconstitution approach: intratracheal challenge of immunocompetent C57BL/6J mice with the re-infected Af-PmV1 strain resulted in pulmonary fungal burdens comparable to those of the parental Af-PmV1-infected Asfu1608 strain and significantly lower than those of the mycovirus-cured Asfu1608r strain (Figure 7D), providing *in vivo* evidence that Af-PmV1 infection directly reduces fungal survival in the lung. Taken together, these data demonstrate that Af-PmV1 infection enhances *A. fumigatus* susceptibility to oxidative- and leukocyte-mediated killing through fungal-intrinsic changes, and that this enhanced killing susceptibility translates to significantly reduced pulmonary fungal burdens *in vivo*.

**Figure 7.**
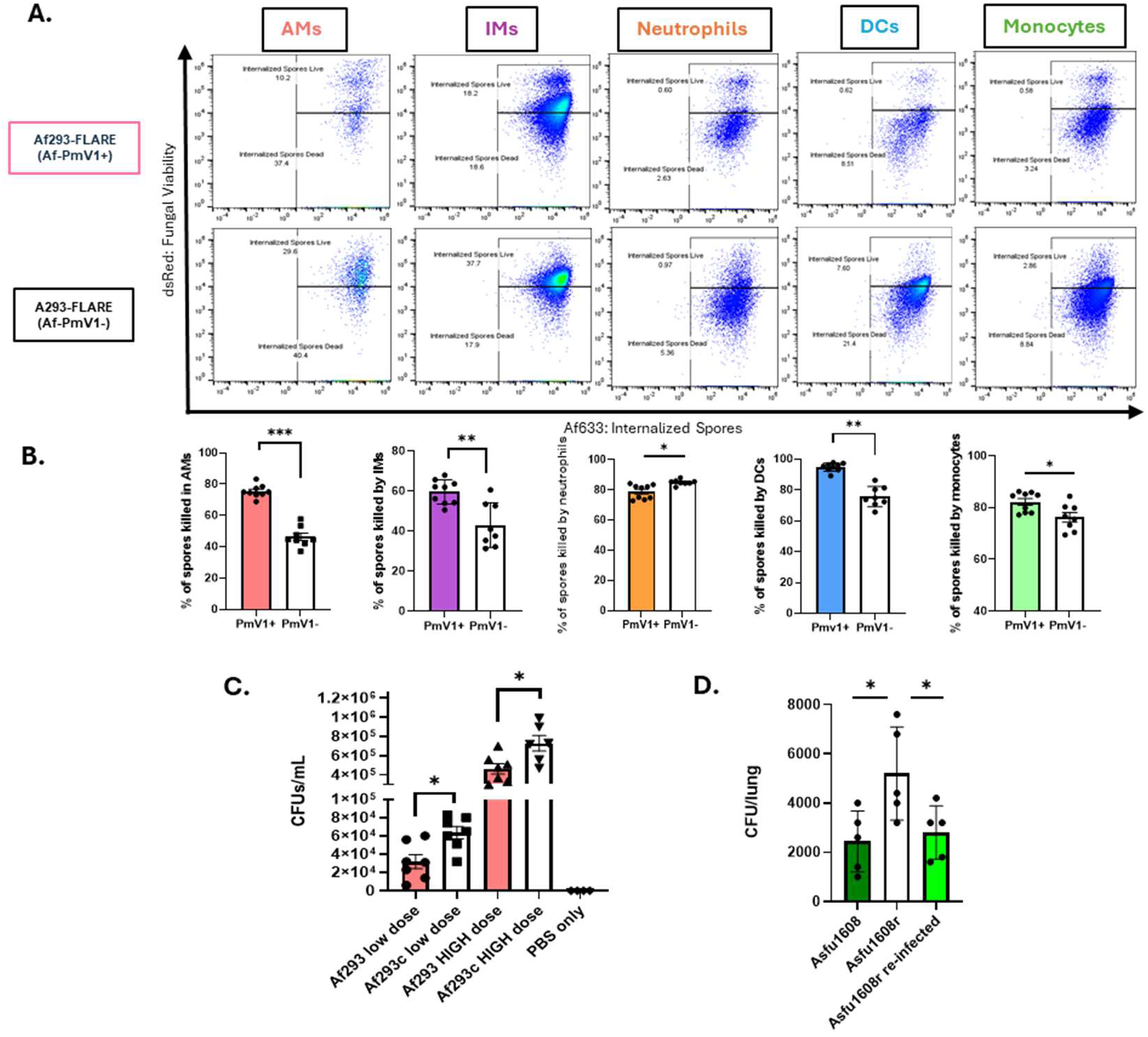
Af-PmV1 infection enhancing *A. fumigatus* killing by alveolar macrophage, but not neutrophils *in vivo* and reduces pulmonary fungal burden. **(A)** Immunocompetent C57BL/6J mice were intratracheally challenged with 4 × 10^7^ spores of the fluorescent reporter strain of AF293 and whole lungs were homogenized in a single-cell suspension. Loss of the dsRed+ spores in various immune cell types were quantified via flow cytometry. Flow plots show data from a single representative mouse. **(B)** Quantification of percent of spores killed in Figure 7A. Each dot represents a single mouse, and the bar represents the mean ± one standard deviation. Statistical significance was determined using a Student’s t-test (\**P* < 0.05; *** *P* < 0.001; **** *P* < 0.0001). **(C)** Immunocompetent C57BL/6J mice were challenged with resting AF293 spores that were either Af-PmV1 infected or Af-PmV1 cured with ribavirin. Mice were given a dose of either 1 × 10^7^ intranasally or 4 × 10^7^ intratracheally. 48 hours post infection, lung CFUs were quantified from total lung homogenate. Each dot represents a single mouse, and the bar represents the mean ± one standard deviation. Statistical significance was determined using a Student’s *t*-test (\**P* < 0.05). **(D)** Immunocompetent C57BL/6J mice were challenged with 4 x 10^7^ resting *Asfu1608* spores intratracheally that were either Af-PmV1 infected or Af-PmV1 cured with ribavirin. Each dot represents a single mouse, and the bar represents the mean ± one standard deviation. Statistical significance was determined using a Student’s *t*-test (\**P* < 0.05).

### Af-PmV1 Infection Alters Virulence in a Chronic ABPA model, but not an acute bronchopneumonia model

Having established that Af-PmV1 infection enhanced *A. fumigatus* susceptibility to H_2_O_2_-mediated killing *in vitro* and to macrophage-mediated killing *in vivo* (Figure 6), we next assessed whether mycoviral infection influenced virulence in murine models of aspergillosis. In an acute bronchopneumonia model, neither AF293/AF293r nor Asfu1608/Asfu1608r strain pairs showed significant differences in mortality (Supplemental Figure 5). This finding was somewhat unexpected, given that mycovirus-cured *A. fumigatus* has been reported to be hypovirulent in a neutropenic model of infection (47), while others have observed no effect on virulence in a corticosteroid model of infection (45).

Given that Af*-*PmV1 infection enhanced susceptibility to H_2_O_2_-mediated and macrophage-mediated killing, we turned to a chronic model of aspergillosis in which *A. fumigatus* conidia persist long-term, driving a Th2- and IgE-dependent disease that recapitulates features of human ABPA (5). Because Asfu1608 was originally isolated from a patient with ABPA (48), this strain was selected for use in the chronic ABPA-like disease model. Immunocompetent C57BL/6J mice were sensitized intranasally with low doses of resting conidia from either Asfu1608 or Asfu1608r on six occasions over two weeks, followed by a final intranasal challenge on day 0 (Figure 8A). Seven days after the final challenge, total serum IgE was quantified and compared against PBS-treated control mice. Both groups exhibited elevated serum IgE; however, mice sensitized with the Af-PmV1 infected Asfu1608 displayed significantly lower serum IgE than mice sensitized with the cured strain Asfu1608c (Figure 8B). Both groups developed significant lung inflammation and fungal persistence, as assessed by lung histology (Figure 8C). These data are consistent with the induction of an ABPA-like disease in mice. Comparing the disease severity between the Asfu1608 and Asfu1608r strains, we observed a significant reduction in serum IgE in the presence of the mycovirus infection (Figure 8B) that was accompanied by decreased in BAL eosinophil and neutrophil counts (Figure 8D-E), indicating that Af-PmV1 infection attenuated the Th2 allergic response. To further characterize the impact of mycoviral infection on fungal persistence, lung sections were assessed by Grocott’s-methenamine silver (GMS) staining and an *Aspergillus-*specific 18S ribosomal qPCR assay. Both strain groups showed evidence of fungal conidia persistence with focal inflammation by GMS staining (Figure 8C); however, mice sensitized with the Af-PmV1-cured Asfu1608r strain exhibited significantly higher pulmonary fungal burdens by qPCR (Figure 8F). Excitingly, when we reexamined the clinical parameters from the original isolation of Asfu1608 (48), we observed that total serum IgE in the patient from which Asfu1608 was the lowest compared to five other patients whose *A. fumigatus* isolate was mycovirus-negative (Supplemental Figure 7). This data is supportive of the role of mycovirus infection in regulating severity of chronic ABPA disease even in humans, but this observation is limited due to having only one mycovirus infection strain. Together, these data demonstrate that Af-PmV1 mycoviral infection limits fungal persistence in a chronic ABPA-like disease model.

**Figure 8.**
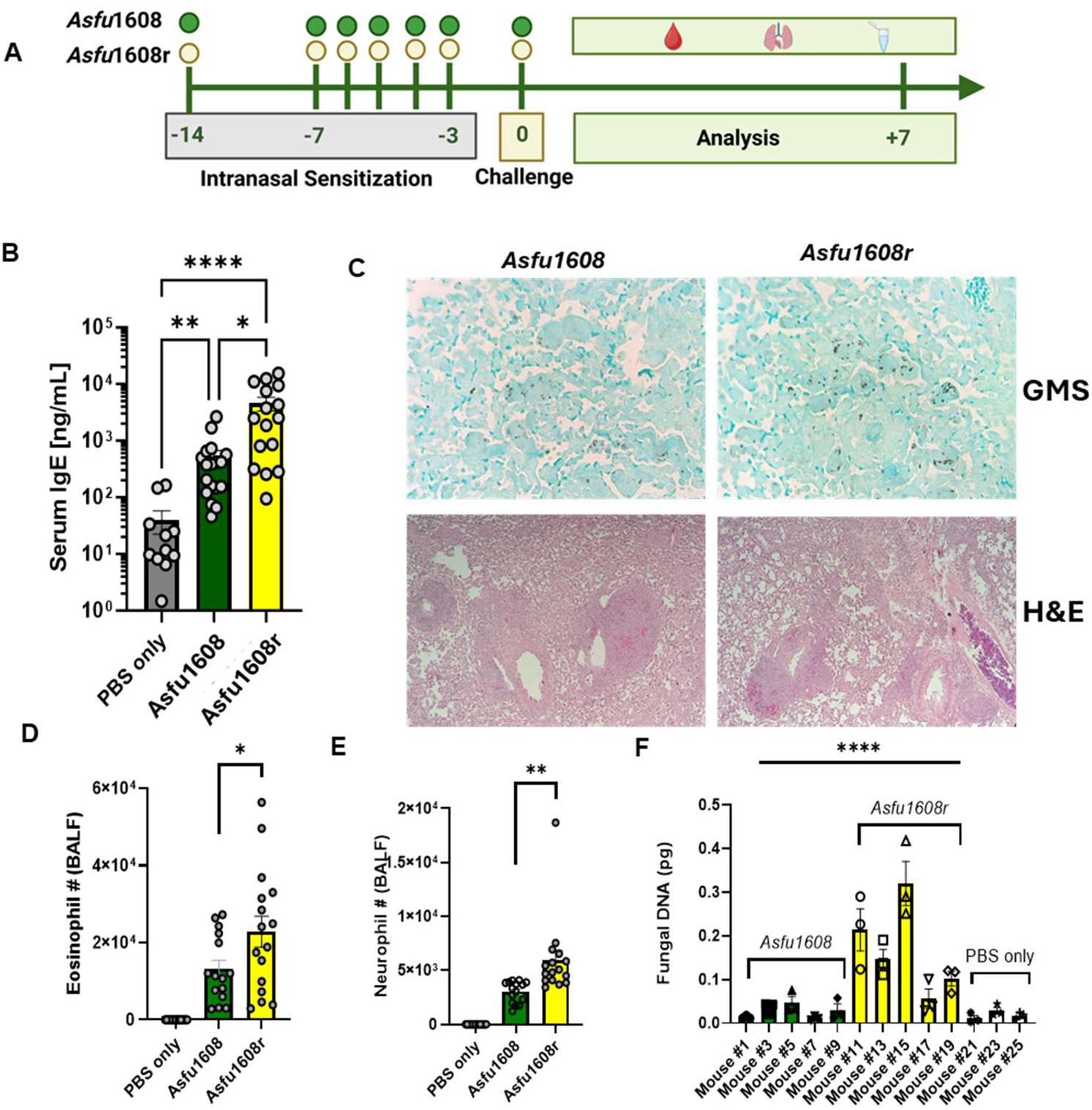
Af-PmV1 mycoviral infection serum IgE, airway eosinophilia, and pulmonary fungal burden in a murine model of allergic bronchopulmonary aspergillosis (ABPA). **(A).** Mice were sensitized intranasally with 1 × 10^7^ resting conidia of Asfu1608 (Af-PmV1-infected) or Asfu1608r (Af-PmV1-cured by ribavirin treatment) for a total of six doses, with the final dose administered on day 0. Mice were euthanized on day 7 post-challenge for collection of bronchoalveolar lavage fluid (BALF) and blood serum by cardiac puncture. **(B)** Total serum IgE was quantified by ELISA. Each symbol represents an individual mouse; bars represent the mean ± SEM. **(C)** Representative Grocott- Gömöri methenamine silver (GMS)- and hematoxylin and eosin (H&E)-stained sections of mouse lungs harvested on day 7 post-challenge. Images were acquired at 40× magnification. **(D-E)** Absolute numbers of eosinophils and neutrophils in BALF on day 7 post-challenge. Each symbol represents an individual mouse; bars represent the mean ± SEM. Statistical significance was determined using a Student’s *t*-test (\**P* < 0.05; ** *P* < 0.01). **(F)** Pulmonary fungal burden was quantified by *Aspergillus-*specific 18S rDNA qPCR from homogenized lung tissue on day 7 post challenge. Each symbol represents an individual mouse; bars represent the mean ± SEM from three independent experiments.

## Discussion

This study demonstrates that Af-PmV1 infection of two *A. fumigatus* strains altered two major dimensions of the host-fungal interaction: (1) enhanced susceptibility to host-relevant reactive oxygen species (ROS) and leukocyte-mediated killing, and (2) augmented type I IFN-dependent inflammatory responses relative to Af-PmV1-cured strains. Collectively, these alterations in the host-fungal interaction reduced the severity of ABPA-like disease in mice sensitized and challenged with mycovirus-infected *A. fumigatus*, highlighting mycoviral infection as a previously unrecognized contributor to chronic aspergillosis outcomes.

Consistent with our findings here that Af-PmV1 infection results in enhanced susceptibility to ROS and leukocyte-mediated killing, prior studies have demonstrated that mycoviral infection broadly enhances fungal susceptibility to host-relevant stresses (13,15,54). Takahashi-Nakaguchi *et al* (47) reported that Af-PmV1-infected AF293 conidia were more susceptible to killing by both H_2_O_2_ and J774A.1 macrophage-like cells than a mycovirus-cured derivative, a finding that corresponded with reduced fungal burden in a cyclophosphamide-induced model of invasive pulmonary aspergillosis. In contrast, Rocha *et al* (50) reported that Af-PmV1-infected AF293 conidia were more resistant to oxidative stress and more virulent in an acute bronchopneumonia model of *A. fumigatus* infection. We initially observed intrinsic differences in hydrogen peroxide killing susceptibility between mycovirus-infected and mycovirus-cured AF293 isolates, which were dependent on the germination state of the conidia at the time of hydrogen peroxide exposure, with resting and pre-swollen conidia exhibiting opposing patterns of susceptibility. This germination state-dependent effect was not observed for the Asfu1608 clinical isolate, in which Af-PmV1 infection enhanced susceptibility to H_2_O_2_ selectively in germinated conidia. The discrepancies across studies likely reflect differences in experimental conditions, including fungal culture and growth conditions, H_2_O_2_ concentration, and/or duration of oxidant exposure. To directly evaluate whether the mycoviral status influences fungal clearance *in vivo*, we employed an acute bronchopneumonia challenge model. Although Af-PmV1 infection was associated with reduced fungal burden for both AF293 and Asfu1608, this reduction was insufficient to produce significant differences in overall virulence in this acute model.

Significant differences in disease parameters emerged when we transitioned to a chronic murine model of allergic bronchopulmonary aspergillosis (ABPA), in which Asfu1608-sensitized mice exhibited marked differences in serum IgE levels, inflammatory cell recruitment, and pulmonary fungal burden depending on the mycoviral infection status of the *A. fumigatus* strain. The reduced disease severity observed in mice challenged with Af-PmV1-infected Asfu1608 likely reflects, at least in part, the fungal-intrinsic increase in conidial susceptibility to ROS-mediated killing conferred by the mycoviral infection, resulting in more efficient early fungal clearance and reduced allergenic stimulus during the sensitization phase. A key unresolved question, however, is whether the augmented type I IFN response driven by Af-PmV1 infection independently contributes to the suppression of IgE seroconversion and ABPA disease severity, or whether reduced fungal antigen load alone is sufficient to explain the observed differences in allergic disease parameters.

Importantly, our murine ABPA findings may be supported by preliminary clinical observations, where total serum IgE levels in patients with ABPA appeared to correlate inversely with the mycoviral status of *A. fumigatus* strain colonizing those patients. Interestingly, initial *Aspergillus*-specific IgE level in ABPA patients found at the time of diagnosis can be used to stratify risks of exacerbation in ABPA patients (51). Although the current patient dataset is insufficient for formal statistical analyses, this observation raises the possibility that mycoviral infection status may be a clinically relevant determinant of ABPA disease severity. The clinical significance of our finding will ultimately depend on the prevalence of Af-PmV1 infection among *A. fumigatus* isolates colonizing patients with ABPA. While the prevalence of mycovirus among clinical *A. fumigatus* isolates remains incompletely characterized, existing data suggest that mycoviruses are detectable in ∼6.6% of *A. fumigatus* strains (52); however, whether the prevalence differs between environmental and clinical isolates, or between isolates from immunocompromised individuals and those from patients with chronic aspergillosis syndromes, remains poorly understood. Thus, a prospective, longitudinal study in which *A. fumigatus* isolates from well-characterized ABPA patient cohorts are routinely screened for mycoviral status and correlated with clinical disease metrics, including total and *Aspergillus*-specific IgE, peripheral eosinophilia, pulmonary function, and radiographic findings, will be an important next step in establishing the translational significance of our findings.

The influence of mycoviral genomes on host innate immune detection of foreign RNA remains largely unexplored. We previously demonstrated that MDA5/MAVS signaling is essential for antifungal immunity against *A. fumigatus* in both mice and humans (20, 21) and that type I and type III IFNs play critical roles in host defense against this pathogen (48). Our earlier work established that dsRNA isolated from *A. fumigatus* is sufficient to activate MDA5 and induce type I IFN expression (20); however, the precise source of immunostimulatory dsRNA within the fungal RNA pool was not identified. Given that MDA5-mediated dsRNA sensing promotes fungal clearance (22) and that the majority of characterized mycoviruses possess dsRNA genomes (15), we reasoned that mycoviral dsRNA may constitute a previously unrecognized pathogen-associated molecular pattern (PAMP) capable of activating MDA5 during *A. fumigatus* infection. In support of this model, both Af-PmV1-infected strains, AF293 and Asfu1608, induced robust type I IFN production in host cells, whereas mycovirus-cured derivatives did not. However, we also observed that the strain CEA10, which appears to lack mycoviral infection, nonetheless elicits a measurable IFN response in mice compared to naïve controls, which was essential for host resistance (20,21). This finding raises the possibility that mycovirus-independent IFN-stimulatory signals exist with *A. fumigatus*, potentially involving alternative innate immune pathways such as Dectin-1-mediated sensing of fungal β-1,3-glucans during both *A. fumigatus* (53) and *Candida albicans* infections (48,49). Additionally, a role for cGAS/STING-mediated IFN responses has been described through the sensing of fungal DNA within extracellular vesicles (56). Together, these observations suggest that the host type I IFN response during *A. fumigatus* infection is driven by multiple, partially redundant innate immune sensing pathways, with mycoviral dsRNA representing a quantitatively dominant, but not exclusive, source of immunostimulatory signal for the antifungal type I IFN response.

The capacity of mycoviral dsRNA to engage host innate immune sensors through trans-kingdom signaling is part of an emerging paradigm in host-pathogen biology. In leishmaniasis, Leishmania RNA virus 1 (LRV1) exacerbates mucocutaneous disease through TLR3-dependent signaling (57). In *Pseudomonas aeruginosa* infection, the filamentous Pf4 bacteriophage triggers TLR3-dependent type I IFN responses that impair bacterial clearance (58) – a mechanism with particular relevance to chronic *P. aeruginosa* pulmonary infections in people with cystic fibrosis (57,58). Within the fungal kingdom, several *Malassezia* species have been shown to harbor totiviruses with dsRNA genomes capable of stimulating type I IFN production from bone marrow-derived macrophages in a TLR3-dependent manner (61), and the dsRNA partitivirus TmV1 has been associated with increased virulence of *Talaromyces marneffei* through aberrant expression of various virulence factors (18) -- though the impact of TmV1 infection on host innate immune responses was not examined in that study and represents an important open question. Extending the relevance of dsRNA sensing to the human airway, TLR3-mediated recongition of *A. fumigatus* dsRNA has been demonstrated in human bronchial epithelial cells (62) and in both human and murine dendritic cells (63,64), suggesting that multiple cell types in the respiratory tract are capable of sensing mycoviral PAMPs. Importantly, the clinical relevance of these innate immune sensing pathways is supported by human genetic data: polymorphisms in *TLR3* (*rs3775296*) and *MAVS* (*rs17857295*) have been associated with increased risk of invasive pulmonary aspergillosis in HSCT patients, and a distinct *TLR3* polymorphism (*rs1879026*) has been associated with ABPA in humans (21,63,65). Collectively, these genetic associations suggest that inter-individual variation in the capacity to sense fungal dsRNA – whether of mycoviral or other origin – may be a significant determinate of aspergillosis susceptibility and disease phenotype in humans and position the MDA5/MAVS and TLR3/TRIF sensing axes as a potential target for host-directed therapeutic strategies in ABPA.

Several limitations of the current study warrant acknowledgment. First, our mechanistic conclusions are based primarily on two naturally Af-PmV1-infected strains, AF293 and Asfu1608, and their respective mycovirus-cured isogenic derivatives. Although the use of isogenic pairs controls for strain background effects, the generalizability of Af-PmV1-mediated immune modulation across the broader *A. fumigatus* population will require confirmation in a larger panel of mycovirus-infected clinical isolates. Second, our *in vivo* experiments were conducted using inbred naïve, specific pathogen-free (SPF) C57BL/6J mice, which may not fully recapitulate the immunological complexity of human ABPA due to the disease developing over years of chronic fungal colonization in the context of underlying structural lung disease, including cystic fibrosis, asthma, and COPD. Third, in our chronic ABPA-like model, Af-PmV1 mycoviral infection limited fungal persistence, likely through fungal-intrinsic mechanisms that reduce conidial resistance to ROS-mediated killing. We did not, however, directly assess the contribution of the enhanced type I IFN-dependent inflammatory response to allergic sensitization, Th2 polarization, or IgE seroconversion in this model – an important limitation that future studies should address. Of note, in murine models of allergic airway disease, both AlvMϕ- and IFN-λ-dependent inflammatory signaling have been shown to be necessary for the induction of Th2/IgE-mediate immunity (59,60). Whether the augmented IFN response driven by Af-PmV1 infection contributes to, or conversely limits, allergic sensitization in ABPA therefore remains an open and important question. Finally, our preliminary human clinical data are cross-sectional and underpowered for formal statistical analysis; a prospective longitudinal study with adequate sample size will be required to establish a causal relationship between mycoviral infection status and ABPA disease severity in human patients. Collectively, these findings reveal a novel role for mycoviral infection in shaping mammalian-fungal interaction in the lung and highlight mycoviruses as previously unrecognized modulators of ABPA immunopathogenesis with potential therapeutic implications. PM

## Acknowledgements

We would like to thank Dr. Kaesi Morelli (current location University of Vermont) for assistance with fungal gDNA prep for whole genome sequencing. This work was supported by grants from the National Institutes of Health: R01 AI139133 (J.J.O), R01 AI191312 (J.J.O.), R35 GM146795 (A.O.), R01 AI130128 (R.A.C.), and R01 AI191067 (R.A.C). A.W.R. was supported by the Dartmouth College Molecular Microbiology & Pathogenesis Program (NIH/NIAID T32 AI007519). A.P.M. was supported by the Dartmouth College Immunology Training Program (NIH/NIAID T32 AI007363). B.S.R. was supported in part by the Dartmouth Cystic Fibrosis Training Program (NIH/NHLBI T32 HL134598). The DartLab Flow Cytometry Core Facility and Dartmouth Health Pathology Shared Resource were supported by the NCI Cancer Center Support Grant P30 CA023108. Core facility support was provided by NIH grant P30 DK117469, NIH grant P20-GM113132 (Dartmouth BioMT COBRE), the CFF Research Development Program, and NCI Cancer Center Support Grant P30 CA023108.

## Authors’ Contribution

AWR, XW, BSR, and AKC conducted the experiments

AWR, BSR generated fungal strains

ARS and KL provided fungal isolates

SMT and AJO generated cell lines

AWR, XW, AKC, RAC and JJO analyzed the experimental data

AWR and JJO wrote the manuscript

