## Supplement Figures for "Trans-Kingdom dsRNA Sensing: *Aspergillus fumigatus* Mycovirus Activates MDA5/MAVS Immunity and Limits Allergic Bronchopulmonary Aspergillosis"

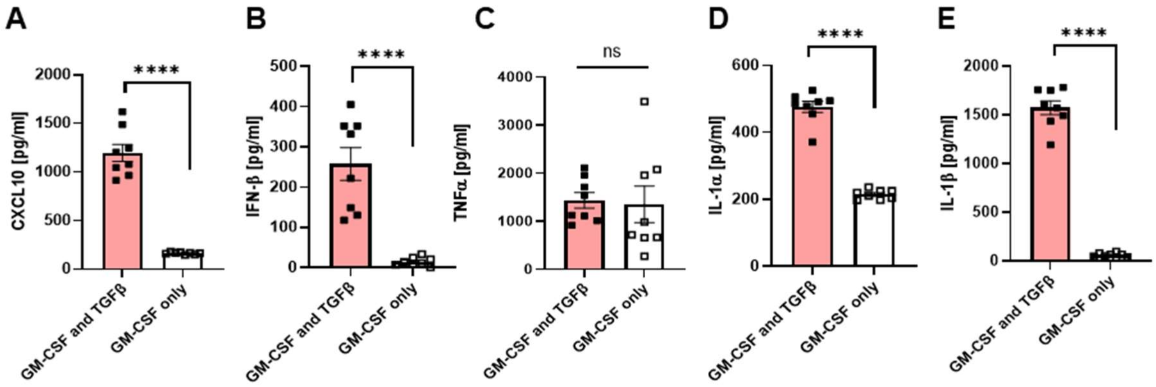


**Supplemental Figure 1. TGFβ-primed alveolar macrophage-like cells preferentially induce a more potent type I IFN response to *A. fumigatus* strain AF293 than *A. fumigatus* strain CEA10. (A-E)** Fetal liver-derived alveolar macrophage-like cells (FLAMs) were cultured with 30 ng/ml recombinant murine GM-CSF, with or without 15 ng/ml recombinant human TGFβ, and challenged overnight with resting conidia of AF293 at an MOI of 10:1. Cytokine concentrations in culture supernatants were quantified by ELISA. Data are pooled from two independent experiments. Each symbol represents a single well; bars represent the mean ± one SD. Statistical significance was determined using a two-way ANOVA (ns = not significant; **** $P < 0.0001$).

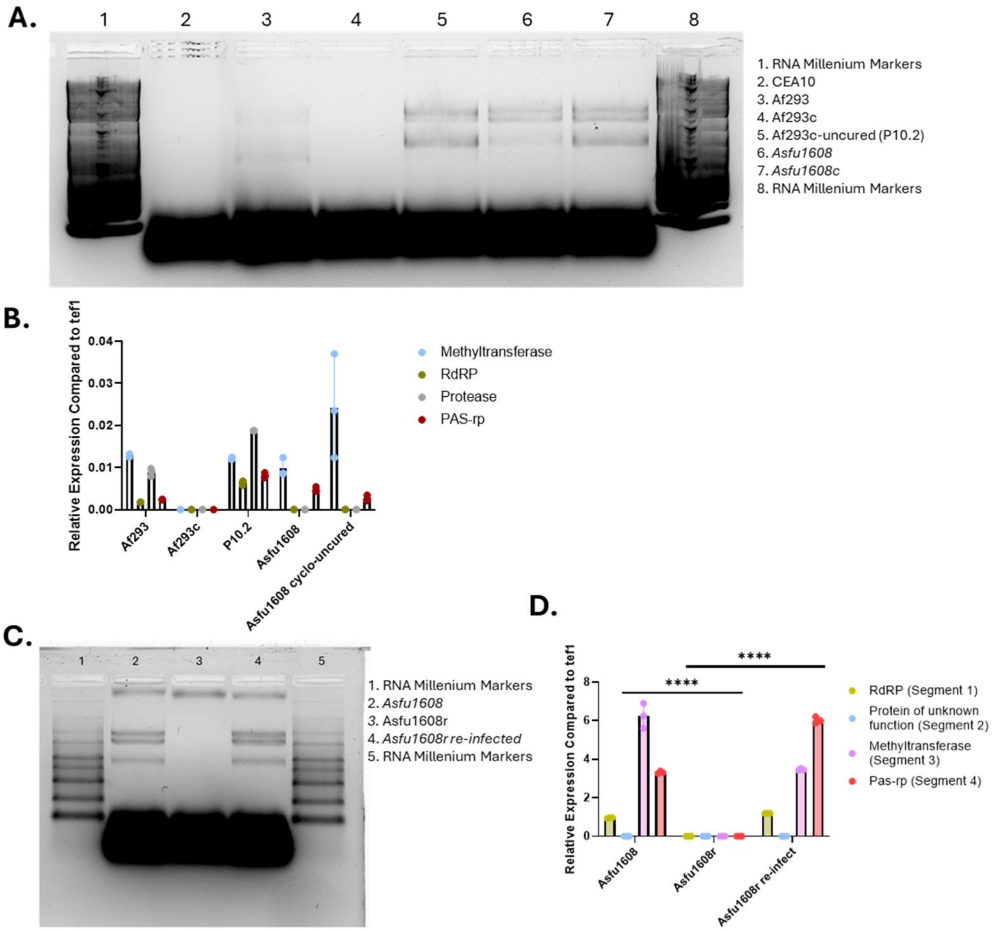


**Supplemental Figure 2. Cycloheximide fails to cure Asfu1608 of Af-PmV1 infection and mycoviral dsRNA transfection successfully re-infects mycovirus-cured Asfu1608r. (A)** 150 µg/ml cycloheximide was used in an attempt to cure the PmV1+ strain Asfu1608 following 10 passages. The presence of the PmV1 methyltransferase gene, RdRp, and PmV1 PAS-RP was screened in various isolates via RT-PCR on the RNA from fungal biofilms and looking for the PCR product on agarose gel. The bands shown in lanes 5, 11, 17, and 23 show the PCR product of Asfu1608 when tested for the presence of *tef1,* the PmV1-methyltransferase gene, RDRP, and Pas-RP respectively. Fungal translation elongation factor gene *tef1* (*Afu1g06390)* used as a control. **(B)** qRT-PCR was performed on AF293, the PmV1-cured isolate of AF293 (AF293c), a PmV1-uncured strain of AF293 that underwent serial passage on cycloheximide plates (P10.2), Asfu1608, and the uncured strain of Asfu1608 that underwent serial passage on cycloheximide plates. **(C)** 10µg of dsRNA from *Asfu1608* was used to re-infect mycovirally-cured *Asfu1608r* fungal protoplasts. The resulting transformants were screened for the re-introduction of Af-PmV1 mycovirus by resolving dsRNA on a 2% NorthernMax™ gel. **(D)** The re-infected strains were screened for the presence of the Af-PmV1 dsRNA segments via qRT-PCR. Each symbol represents a technical replicate; bars represent the mean ± one SD. Statistical significance was determined using a two-way ANOVA (**** $P < 0.0001$).

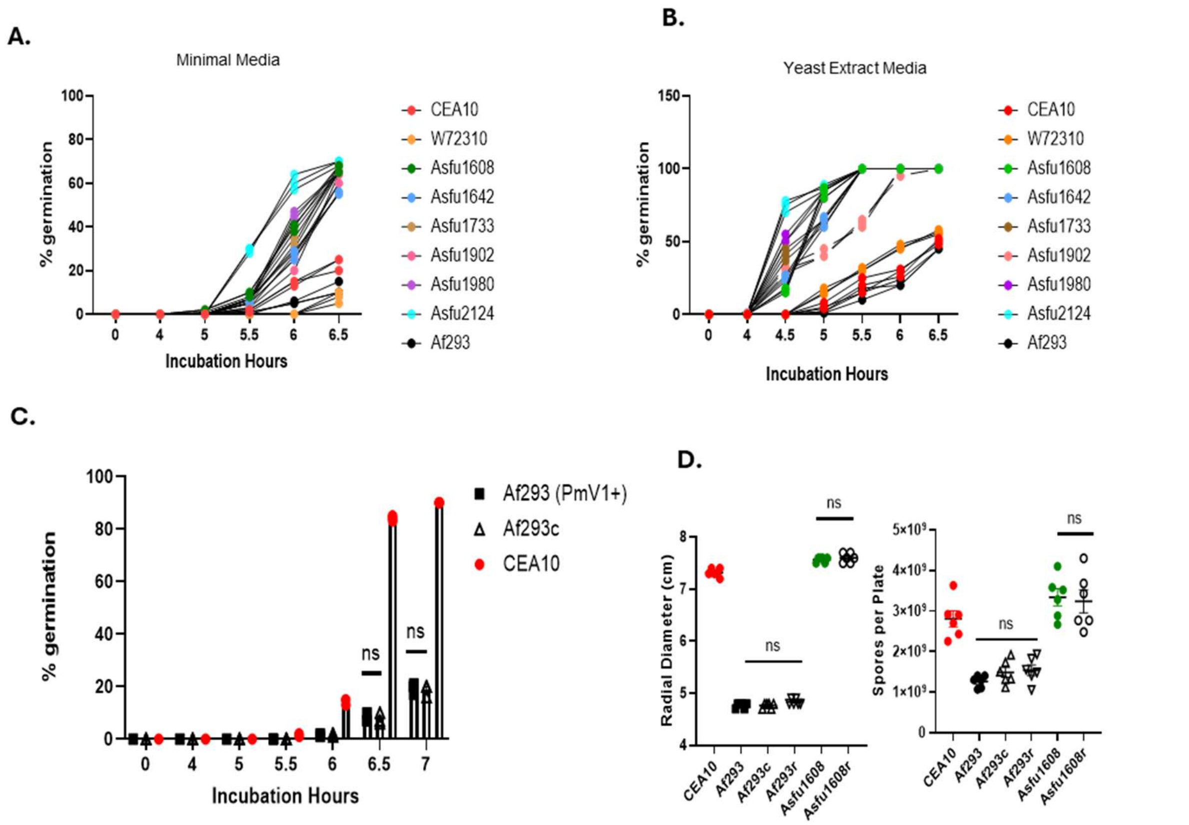


**Supplemental Figure 3. Af-PmV1 infection does not alter *A. fumigatus* radial growth, sporulation, or germination rates. (A-B)** Various strains were grown in either L-GMM or liquid yeast extract medium at $5 \times 10^6$ conidia/ml with shaking at 37°C in normoxic conditions. Percent germination was monitored over time. **(C)** Germination rates of the Af-PmV1+ and isogenic cured strain of AF293 were grown in L-GMM. Each symbol represents an independent culture flask; bars represent the mean ± SD. Statistical significance was determined using a two-way ANOVA (**** $P < 0.0001$). **(D)** Radial growth of PmV1+ and cured strains grown on GMM plates for five days. Sporulation after five days of PmV1+ and cured strains grown on GMM plates. Each shape represents an independent plate, and statistical significance was determined using a two-way ANOVA (NS = non-significant).

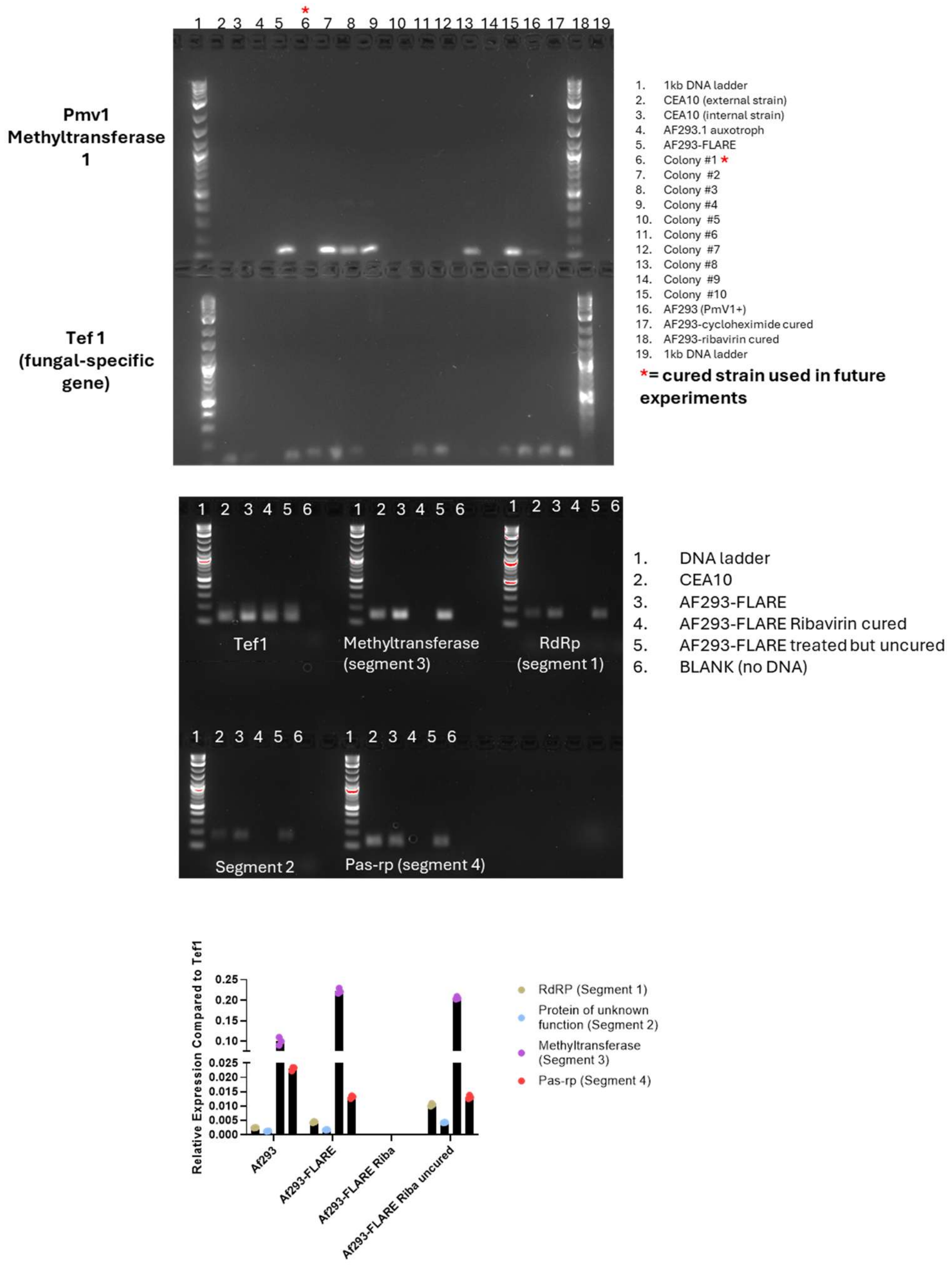


**Supplemental Figure 4. Ribavirin treatment successfully cures the fluorescent AF293-FLARE reporter strain of Af-PmV1 infection.** The presence of the PmV1 methyltransferase gene was screened in various isolates via RT-PCR on the RNA from fungal biofilms and looking for the PCR product on agarose gel. The isolate in lane 6 (red asterisk, Colony #1) was used as the cured strain in all future experiments. Fungal translation elongation factor gene *tef1* (*Afu1g06390)* used as a fungal-specific control gene.

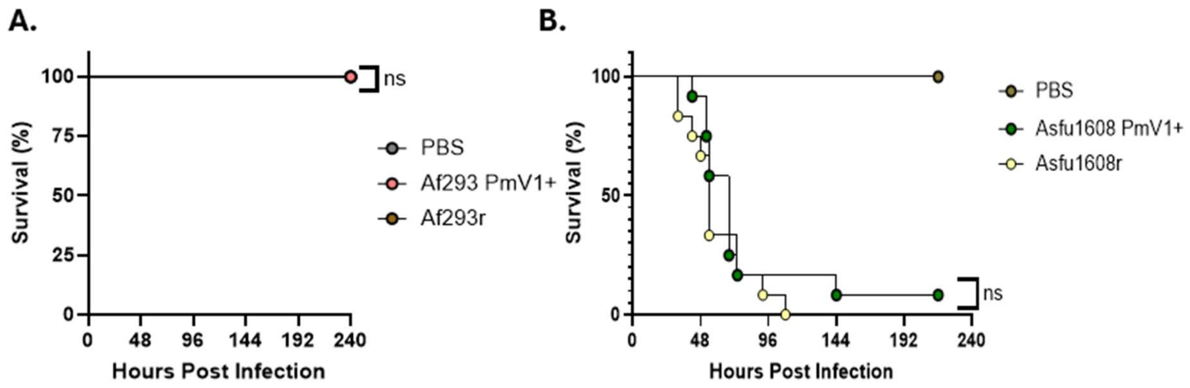


**Supplemental Figure 5. Af-PmV1 infection does not influence mortality in an acute bronchopneumonia model. (A)** Neither AF293 or AF293r strains or **(B)** Asfu1608 and Asfu1608r strains showed significant differences in mortality. Mice were monitored for 10 days for health and survival. Data are representative of 2 independent experiments containing 8-12 mice per group. Mouse survival data were analyzed with the Mantel-Cox log rank test (NS = non-significant).

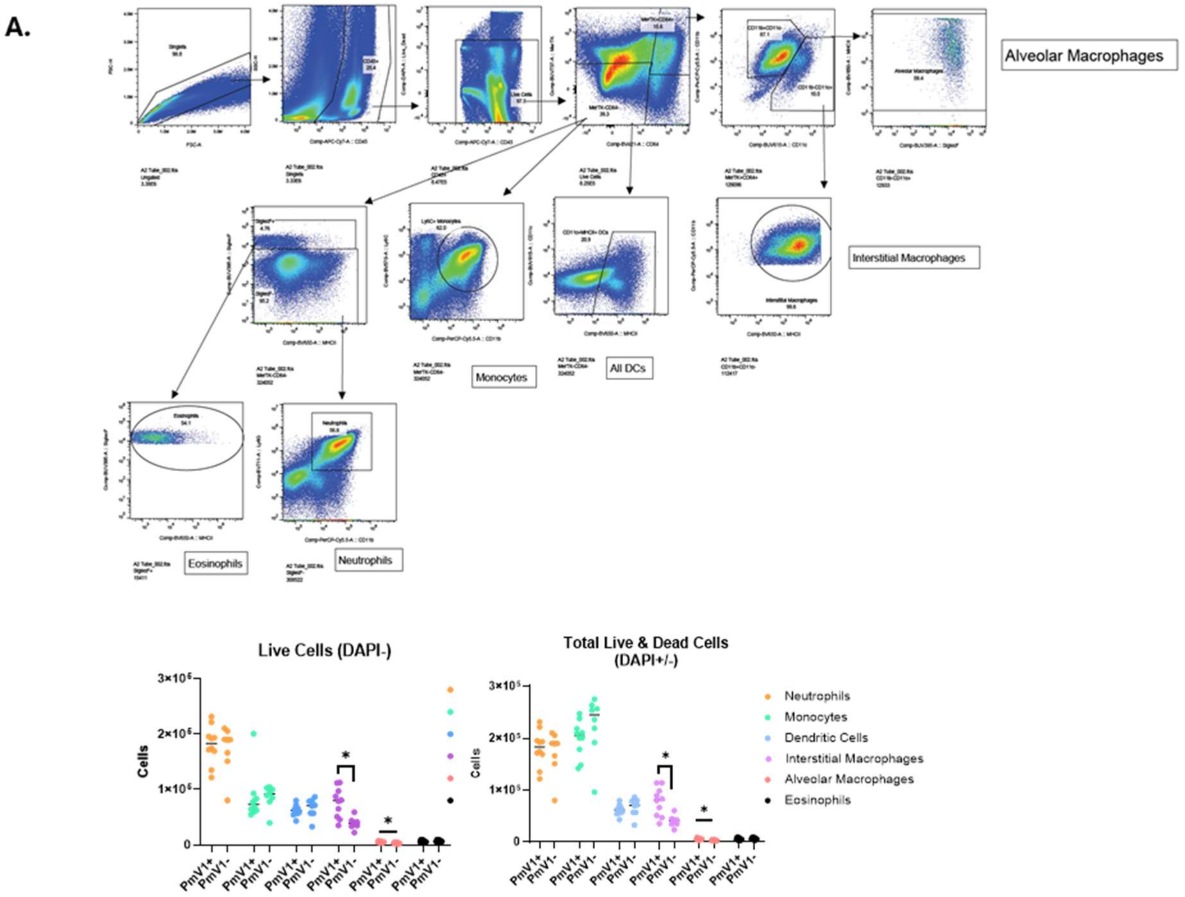


**Supplemental Figure 6. Flow cytometry gating strategy for identification of lung myeloid cell populations from in AF293-FLARE-infected C57BL/6J mice. (A)** Gating strategy of C57BL/6J mice infected with AF293-FLARE strain of Aspergillus. **(B)** Cell counts of whole-lung homogenate in mice infected with AF293-FLARE strain. Each symbol represents a mouse; bars represent the mean ± SD Statistical significance was determined using a Student's *t*-test (NS = non-significant; ** $P < 0.01$).

**A.**

| Strain ID | IgE (g/L) | IgG (g/L) | *Af* precipitin | Mycoviral Status |
|---|---|---|---|---|
| Asfu1608 | 170 | 10.7 | + | + |
| Asfu1642 | 514 | 20.5 | + | - |
| Asfu1733 | 2024 | 16 | + | - |
| Asfu1902 | 404 | 6.16 | + | - |
| Asfu1980 | 600 | 6.81 | + | - |
| Asfu2124 | 671 | 11.5 | + | - |

**B.**

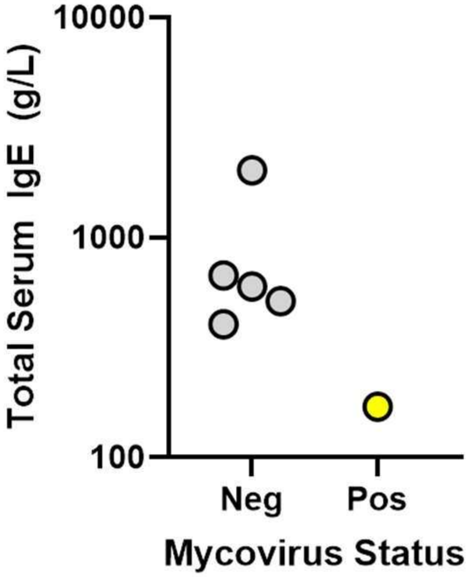


**Supplemental Figure 7. A mycovirus-positive *A. fumigatus* clinical isolate is associated with lower serum IgE levels in patients with allergic bronchopulmonary aspergillosis (ABPA). (A)** Table of *A. fumigatus* isolated from the Vermeulen *et al.* (48) study with the representative clinical data from each of those patients. Mycoviral infection status was determined in our current study -- see Figure 3 for dsRNA gels. **(B)** Graphical depiction of total serum IgE levels form ABPA patients separated by mycoviral infection status. Each symbol represents a single individual.

**Supplemental Table 1**: Strains used and generated for this work.

| Strain | PmV1-infected | Drug Exposed To | Source |
|---|---|---|---|
| CEA10 | Negative | N/A | Girardin et al., 1993 (68) |
| AF293 | **Positive** | N/A | Pain et al., 2004 (69) |
| AF293c | Negative | Cycloheximide | This study |
| AF293r | Negative | Ribavirin | This study |
| P10.2 | **Positive** | Cycloheximide | This study |
| AF293rib-uncured | **Positive** | Ribavirin | This study |
| W73210 | Negative | N/A | Jones et al., 2021(5) |
| Asfu1608 | **Positive** | N/A | Vermeulen et al., 2015 (48) |
| Asfu1608r | Negative | Ribavirin | This study |
| Asfu1608rib-uncured | **Positive** | Ribavirin | This study |
| Asfu1608cyclo-uncured | **Positive** | Cycloheximide | This study |
| Asfu1642 | Negative | N/A | Vermeulen et al., 2015 (48) |
| Asfu1733 | Negative | N/A | Vermeulen et al., 2015 (48) |
| Asfu1902 | Negative | N/A | Vermeulen et al., 2015 (48) |
| Asfu1980 | Negative | N/A | Vermeulen et al., 2015 (48) |
| Asfu2124 | Negative | N/A | Vermeulen et al., 2015 (48) |
| AF293-FLARE | **Positive** | N/A | Jhingran et al., 2012 (36) |
| AF293-FLARE (PmV1-) | Negative | Ribavirin | This study |

**<u>Supplemental Table 2</u>:** PmV1 and *Aspergillus* Primers Used

| **Name** | **Direction** | **Sequence (5'->3')** |
|---|---|---|
| Tef1 (*Afu1g06390)* | Forward | AAGTATGAGGTCACTGTCATCGAT |
| Tef1 (*Afu1g06390)* | Reverse | ATACCAGCCTCGAACTCACC |
| Methyl transferase 1 | Forward | CGACTACTGCTCTGCCATCC |
| Methyl transferase 1 | Reverse | AACAACTCGGGCTCTACTGC |
| Methyl transferase 2 | Forward | GATGCCATGTACCCCGTCAA |
| Methyl transferase 2 | Reverse | GACCCCATGCTCACCATCAT |
| Methyl transferase 3 | Forward | TCGCGTACAATGCTGAGGTT |
| Methyl transferase 3 | Reverse | TGGAAGAACTGCCACACCTG |
| RDRP1 | Forward | CCATGTCCTCCTCCCTCCTT |
| RDRP1 | Reverse | TCGACGATCTTGGCTGCTTT |
| Protease 1 | Forward | CGTTACGACCCTGCCAAGAT |
| Protease 1 | Reverse | AGGAAACGACGTCCTGGTTC |
| PAS-rp 1 | Forward | CATTCACGCGTGGTCCTTTG |
| PAS-rp 1 | Reverse | CGTCCTTAGGATCCGTGTGG |
| 18S rDNA | Forward | GGCCCTTAAATAGCCCGGT |
| 18S rDNA | Reverse | TGAGCCGATAGTCCCCCTAA |
| 18s rDNA | TaqmanProbe | AGCCAGCGGCCCGCAAATG |

**Supplemental Table 3:** Antibody clones experimentally used.

| Antigen | Clone/Cat. No | Company | Conjugate |
|---|---|---|---|
| CD45 | 30-F11 | Biolegend | APC-Cy7 |
| CD64 | X54-5/7.1 | Biolegend | BV421 |
| CD11b | M1/70 | Biolegend | PerCPCy5.5 |
| SiglecF | E50-2440 | BD Biosciences | BUV395 |
| Ly6G | 1A8 | Biolegend | BV711 |
| MHCII | M5/114.15.2 | BD Biosciences | BV650 |
| MerTK | 367-5751-82 | ThermoFisher | BUV737 |
| CD103 | 2E7 | Biolegend | Af488 |
| CD11c | N418 | ThermoFisher | BUV615 |
| Streptavidin-Af633 | S21375 | ThermoFisher | Af633 |
| DAPI | 422801 | Biolegend | DAPI |

**A.**

| Unique SNPs in Af293c | Unique SNPs in P10.2 | Shared SNPs in cycloheximide treatment | Unique SNPs in Af293r | Unique SNPs in Af293rib-uncured | Shared SNPs in ribavirin treatment |
|---|---|---|---|---|---|
| Afu2g04430 | Afu1g01570 | Afu1g00480 | Afu2g00110 | Afu1g00480 | Afu3g10200 |
| Afu3g05830 | Afu2g04420 | Afu1g09300 | Afu3g12540 | Afu1g01570 | Afu8g04070 |
| Afu3g07870 | Afu4g09260 | Afu2g17600 | Afu4g00820 | Afu1g09300 | |
| Afu3g08490 | Afu5g10520 | Afu3g10200 | Afu6g03610 | Afu2g17600 | |
| Afu4g00970 | Afu7g06330 | | Afu7g01360 | Afu3g05830 | |
| Afu4g04612 | | | Afu8g04020 | Afu3g08490 | |
| | | | Afu8g07400 | Afu3g14700 | |
| | | | | Afu4g00770 | |
| | | | | Afu4g04740 | |
| | | | | Afu7g06330 | |
| | | | | AfuMt00001 | |

**B.**

| Unique SNPs in Asfu1608r | Unique SNPs in Asfu1608rib-uncured | Shared SNPs in ribavirin treatment |
|---|---|---|
| Afu2g11730 | Afu4g00820 | Afu2g00860 |
| Afu3g11180 | Afu6g04570 | Afu2g00870 |
| Afu3g13860 | Afu6g06900 | Afu3g12410 |
| Afu1g11740 | Afu3g02650 | Afu1g09300 |
| Afu2g04490 | Afu5g00100 | Afu2g00110 |
| Afu3g06810 | Afu6g10110 | Afu5g14950 |
| Afu3g10430 | | Afu7g00090 |
| Afu3g13610 | | |
| Afu3g14060 | | |

**C.**

| AF293 Exposed to Cycloheximide | Af293 Exposed to Ribavirin | Asfu1608 Exposed to Ribavirin | % of SNPs shared among each condition |
|---|---|---|---|
| 36.5% | 11.1% | 46.6% | 31.4% |

**Supplemental Table 4. Single-nucleotide polymorphism (SNP) analysis of fungal genome following antiviral drug exposure during the mycoviral curing process. (A)** Genomic DNA sequencing was performed on the strain PmV1-infected AF293 WT strain, as well as the cycloheximide-cured (AF293c), cycloheximide-treated but uncured (P10.2), ribavirin-cured strain (AF293r), and the ribavirin-treated but uncured strain (AF293rib-uncured). DNA mutations unique to each cured and uncured strain were compared to a reference genome. **(B)** The PmV1-infected strain Asfu1608 as well at the ribavirin-cured strain (Asfu1608r) and ribavirin-treated but uncured strain (Asfu1608rib-uncured) were sequenced. DNA mutations unique to the ribavirin-cured and uncured strain were compared to the reference genome.**(C)** The percentage of SNPs shared among both cured and uncured strains following exposure to the same anti-viral drug.
